# CRISPR screens in the context of immune selection identify *CHD1* and *MAP3K7* as mediators of cancer immunotherapy resistance

**DOI:** 10.1101/2025.04.15.648737

**Authors:** Alex Watterson, Gabriele Picco, Vivien Veninga, Youhani Samarakoon, Chiara M. Cattaneo, Sara F. Vieira, Emre Karakoc, Shriram Bhosle, Thomas W. Battaglia, Sarah Consonni, Timotheus Y. F. Halim, Emile E. Voest, Mathew J. Garnett, Matthew A. Coelho

## Abstract

Cancer immunotherapy is only effective in a subset of patients, highlighting the need for effective biomarkers and combination therapies. Here we systematically identify genetic determinants of cancer cell sensitivity to anti-tumor immunity by performing whole-genome CRISPR/Cas9 knock-out screens in autologous tumoroid-T cell co-cultures, isogenic cancer cell models deficient in interferon signaling, and in the context of four cytokines. We discover that loss of *CHD1* and *MAP3K7* potentiates the transcriptional response to IFN-γ, thereby creating an acquired vulnerability through sensitizing cancer cells to tumor-reactive T cells. Immune checkpoint blockade was more effective in a syngeneic mouse model of melanoma deficient in *Chd1* and *Map3k7* and was associated with elevated intra-tumoral CD8^+^ T cell numbers and activation. *CHD1* and *MAP3K7* are recurrently mutated in cancer and reduced expression in tumors correlates with response to immune checkpoint inhibitors in patients, nominating these genes as potential biomarkers of immunotherapy response.

## Introduction

Immune checkpoint blockade (ICB) is an effective treatment for many cancer types, such as melanoma and cancers with microsatellite instability (MSI)^1^, however response rates in other solid tumors are below 35 %^2^. Understanding the genetic determinants of ICB response would help to identify effective combination therapies^3^ and stratify individuals that benefit from treatment, sparing others treatment-related toxicities^4^.

For patients who initially respond to ICB, acquired resistance can emerge through various mechanisms^5^, including *HLA* or *B2M* mutation^6,7^ or disabling the IFN-γ pathway through mutations in *JAK1*/2^8,9^, highlighting the importance of tumor-cell-intrinsic signaling in immune evasion. CRISPR/Cas9 knock-out (KO) screens have identified genetic modulators of IFN-γ response and sensitivity to T cells^10–12^. However, these studies typically cannot discriminate independent effects of different cytokines and deconvolute pathway-specific biology^13,14^, rely on overexpression of artificial antigens^15^, or focus on mouse model systems that use highly immunogenic cell lines^13,14^.

Loss of tumor suppressor genes drive cancer progression and therapy resistance^16^, but can also have important non-cell autonomous effects^17^ and result in acquired vulnerabilities^18,19^. *CHD1* (chromodomain helicase DNA binding protein 1) and *MAP3K7* (encoding transforming growth factor β activated kinase 1; TAK1) have been proposed as tumor suppressor genes^20–22^ due to their recurrent mutation in prostate cancers^23,24^ and loss at lower frequencies in other cancer types^25,26^. In prostate cancer, co-deletion is associated with aggressive disease and therapy resistance^27–29^. Loss of *CHD1*, a chromatin remodeling enzyme^30^, is associated with extensive changes in the tumor microenvironment (TME) in mice through reduced NF-*κ*B signaling^31,32^. *MAP3K7* is found within a commonly deleted genomic region in prostate cancer (chromosome 6q15),^29^ and encodes a kinase involved in activating NF-*κ*B and TGF-β signaling^33–35^. The effects of *MAP3K7* loss on the TME and ICB response remain largely unexplored.

Here we investigate the genetic landscape of sensitivity to autologous, human T cells and cytokines found in the TME in four cancer cell models, generating a genome-scale map of context-specific dependencies. Integrative analysis reveals that *CHD1* and *MAP3K7* loss additively sensitizes cancer cells to IFN-γ and anti-tumor T cells and reduced expression in tumors is associated with patient response to ICB.

## Results

### Functional genomics identifies genetic modulators of cytokine response

To systematically investigate the genetic determinants of cytokine response in cancer cells, we performed whole-genome CRISPR/Cas9 KO screens (Methods^36^) in the presence of cytokines found in the TME (Fig 1A). Firstly, we tested IFN-γ, IFN-β, IL-6, which affect anti-tumor immunity, and depend on JAK1 or JAK2 signaling^37^ (Fig. 1A). We selected cell models from cancer types where immunotherapy is used clinically, such as melanoma^38^ (A375 and SK-MEL-2) and colorectal cancer^39^ (HT-29) and generated two independent isogenic *JAK1* or *JAK2* KO clones for each cell model (Fig. S1A-D) to investigate JAK1 or JAK2-specific signaling dependencies (Fig. 1B). The IFN-γ and IL-6 receptors engage both JAK1 and JAK2^40,41^, whereas the IFN-β receptor engages JAK1 and TYK2^41^, consistent disparate cytotoxic responses to IFN-β in *JAK1* and *JAK2* KO SK-MEL-2 clones (Fig. 1B and S1E). Replicate correlation for independent CRISPR/Cas9 KO screens in the absence of cytokine stimulation was high, verifying screen quality (*r* = 0.82-0.91; Fig. S1F-G). Variability arose in the IL-6 conditions, which had minimal impact on cell growth despite stimulation of signaling (Fig. S1A-D). SK-MEL-2 *JAK1* KO screens highlighted known IFN-β biology including sensitizer (*ADAR*^42^, *PTPN2*^10^ KO) and resistance hits (*TYK2*, *STAT2* and *IFNAR1/2* KO). These were absent in *JAK2* KO screens (Fig. S2A), consistent with the contribution of JAK1, but not JAK2, in IFN-β signaling.

**Figure 1.**
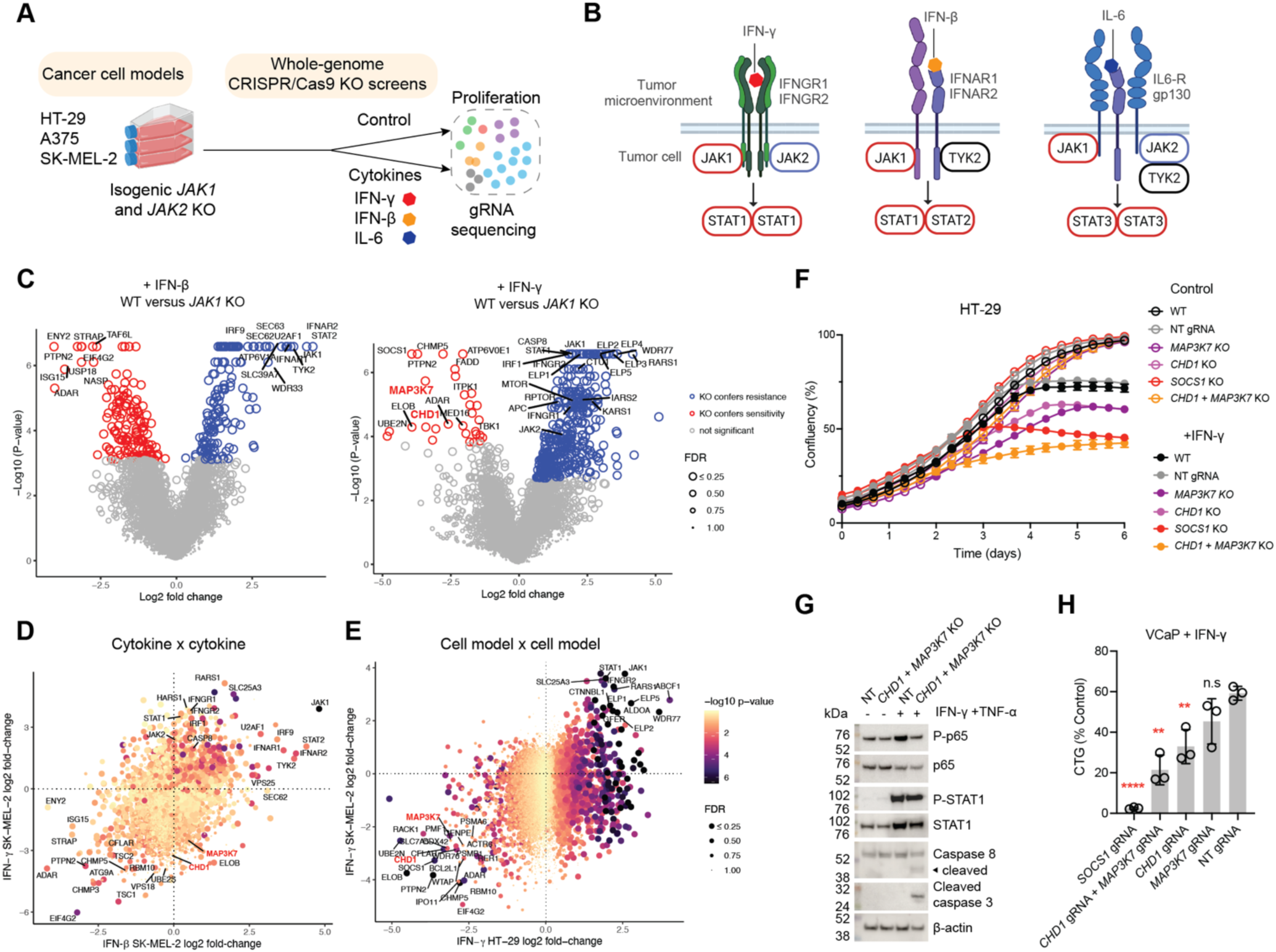
Whole-genome CRISPR screens define the genetic determinants of cytokine sensitivity in cancer cells. **A)** Overview of CRISPR/Cas9 screens to identify modulators of IFN-γ, IFN-β, and IL-6 cytokine response in three cancer cell models. **B)** Overview of cytokine signaling pathways and differential dependencies on JAK1/2. **C)** Gene-level volcano plots of CRISPR/Cas9 KO screens comparing wild-type (WT) and *JAK1* KO SK-MEL-2 cells treated with IFN-β (400 U/ml) (left panel), or HT-29 cells treated with IFN-γ (500 U/mL) (right panel). Data represent the average of two independent screens and significant hits are highlighted (*P*-adjusted <0.05, FDR <0.1). **D)** Shared and private modulators of IFN-γ and IFN-β response in SK-MEL-2 cells. Scatter plots comparing the log2 fold-change of WT control versus IFN-γ or IFN-β-treated *JAK1* KO clones. Data represents the average of two independent screens and are representative of two independent *JAK1* KO clones. IFN-γ or IFN-β screens FDR and *P*-values are indicated for the cytokine with the largest effect size. **E)** CRISPR screens identify genes conferring resistance or sensitivity to IFN-γ in SK-MEL-2 and HT-29 cancer cell models. Scatter plots comparing log 2-fold-change values for WT control versus IFN-γ-treated conditions. *P* adjusted values, and FDR values for HT-29 screens are indicated. Data represent the average of two independent screens. **F)** *CHD1* and *MAP3K7* KO additively sensitize cancer cells to IFN-γ. Cell growth Incucyte curves from *CHD1, MAP3K7, CHD1* and *MAP3K7* dKO, and *SOCS1* KO HT-29 ± IFN-γ (500 U/mL). Data represent the mean ± S.D of three biological replicates and are representative of two independent experiments. Data are representative of two different gRNAs targeting *CHD1* or *MAP3K7*. **G)** *CHD1* and *MAP3K7* KO primes cancer cells for apoptosis in response to IFN-γ and TNF-*⍺*. Western blotting of HT-29 NT gRNA control cells or *CHD1* and *MAP3K7* dKO cells ± IFN-γ (500 U/mL) and TNF-*⍺* 100 (100 ng/ml) for 8 hours. Data are representative of two independent experiments. **H)** Sensitivity of *CHD1* and *MAP3K7* KO prostate cancer cells to IFN-γ. VCaP cells were grown ± IFN-γ (500 U/mL) and cell viability was measured using Cell Titer-Glo (CTG). Data was normalized to untreated controls and represent the mean ± S.D of three independent experiments performed in technical triplicate. Two-tailed, unpaired Student’s t-test; ** *P* <0.01; **** *P* <0.0001; not significant, n.s.

To investigate variation in genetic mediators of response between different cytokines, we compared dependencies in IFN-γ and IFN-β screens (Fig. 1C-D). *ADAR* and *PTNP2* KO were sensitizing, and *JAK1* KO conferred resistance to both cytokines, whereas KO of the genes encoding the receptors for IFN-γ or IFN-β (*IFNGR1/2* and *IFNAR1/2*) provided cytokine-specific resistance. Genes involved in vesicular trafficking and mTOR signaling were sensitizing hits in both cytokine contexts, including *TSC1*, *TSC2*, *CHMP3* and *CHMP5* (Fig. S2B).

Comparison of IFN-γ response modulating genes across cancer cell models identified KO of *JAK1*, *STAT1*, *IRF1* and *CASP8* as conferring resistance in all three cell models (Fig. S2C). KO of *CFLAR*^43^ was universally sensitizing to IFN-γ, consistent with a caspase 8-mediated mechanism of apoptosis^44^. *MTOR* and *RPTOR* were shared resistance hits, implicating cell-intrinsic mTOR signaling in cancer cell IFN-γ response. Other shared resistance hits pertained to tRNA processing (e.g. *RARS1*) and amino acid transport (e.g. *SLC25A3*) (Fig. S2D). These genes are thought to relate to mTOR signaling through modulating cellular amino acid levels^45,46^. IFN-γ sensitizing hits shared across three cancer cell models (*n* = 8) included *TSC1*, *SLC7A5*, *RAB7A* and *CHMP5*, relating to mTOR and vesicular trafficking. Moreover, gene ontology and pathway analysis of shared sensitizing and resistance hits across cancer cell models revealed significant enrichment in tRNA processing and translation (Fig. S2E).

Through screening 15 cell models (WT, *JAK1* KO, and *JAK2* KO) treated with three cytokines, we mapped context-specific genetic dependencies from 138 whole-genome CRISPR/Cas9 KO screen samples (Table S1). Collectively, these data outline both conserved and specific genetic modulators of cytokine signaling in cancer cells and highlight the importance of the mTOR pathway in mediating cancer cell-intrinsic cytokine responses^13,41,47^.

### *CHD1* and *MAP3K7* loss sensitizes cancer cells to IFN-γ

To discover genes that could serve as biomarkers of cancer response to inflammatory cytokines using this combined dataset, we focused on IFN-γ-sensitizing hits with minimal impact on cell fitness in the absence of IFN-γ, such as *PTPN2* and *SOCS1* (Fig. S2F). Two such hits were *CHD1* and *MAP3K7* (Fig. 1E, Fig. S2F-G and Fig. S3A). *CHD1* and *MAP3K7* are co-deleted in prostate cancers and are on separate chromosomes^28^, implying a cooperative mechanism. Consistently, analysis of 51 studies from The Cancer Genome Atlas^48^ (TCGA) and literature^27,49–69^ highlighted prostate cancer as having the highest incidence of loss; 1.2-16.9 % of tumor samples had deletion of one gene, and 0.7-3.9 % had co-deletions of *CHD1* and *MAP3K7* (Fig. S3B). Other cancer types with recurrent co-deletions included thyroid cancer (4.7 %), melanoma (0.2-3.7 %) and meningioma (0.8 %).

To validate our findings from genome-wide CRISPR/Cas9 KO screens, we performed arrayed KO experiments. *CHD1* or *MAP3K7* KO (Fig. S3C) did not alter cell proliferation rate in HT-29 but sensitized to IFN-γ (Fig. 1F). To model co-deletion, we knocked-out both genes together (double KO, dKO). This led to more profound sensitization to IFN-γ than either single gene KO, indicating an additive effect (Fig. 1F). Single KO and dKO HT-29 cells had similar levels of JAK-STAT activation to WT cells in response to IFN-γ or IFN-γ and TNF-*⍺* (P-STAT1; Fig. 1G and Fig. S3D), but reduced NF-*κ*B activity (P-p65) and increased induction of apoptosis (cleaved-caspase 3 and 8; Fig. 1G and Fig. S3D). Takinib, a MAP3K7 (TAK1) inhibitor^70^ phenocopied *MAP3K7* loss on IFN-γ sensitivity, specifically in the context of *CHD1* KO (Fig. S3E). Furthermore, in the prostate cancer model VCaP, dKO cells had increased sensitivity to IFN-γ (Fig. 1H and Fig. S3F). Taken together, these data highlight *CHD1* and *MAP3K7* loss as a cytokine-dependent vulnerability in cancer cells.

### Autologous tumoroid-T cell co-culture CRISPR screens identify modulators of cancer cell sensitivity to tumor-reactive T cells

Stimulation with individual cytokines facilitates the investigation of specific signaling pathways involved in anti-tumor immunity but cannot recapitulate the combination of factors present at the synapse between T cells and tumor cells. To investigate the genetic determinants of tumor cell sensitivity to tumor-reactive T cells more directly, we established a co-culture system for CRISPR/Cas9 KO screening comprising primary tumoroids derived from an MSI colorectal cancer (CRC) (CRC-9) and autologous, tumor-reactive T cells (predominantly CD8^+^ T cells^71^) from peripheral blood mononuclear cells (Methods^12,72–74^). To achieve genome-scale screening with primary material, we used a condensed and efficiency-optimized gRNA library - MinLibCas9^36^. In addition, we screened CRC-9 tumoroids in the presence of IFN-γ and TNF-*⍺*, cytokines involved in T cell-mediated killing, to deconvolute the contributions of these factors in cancer cell death^75^ (Fig. 2A).

**Figure 2.**
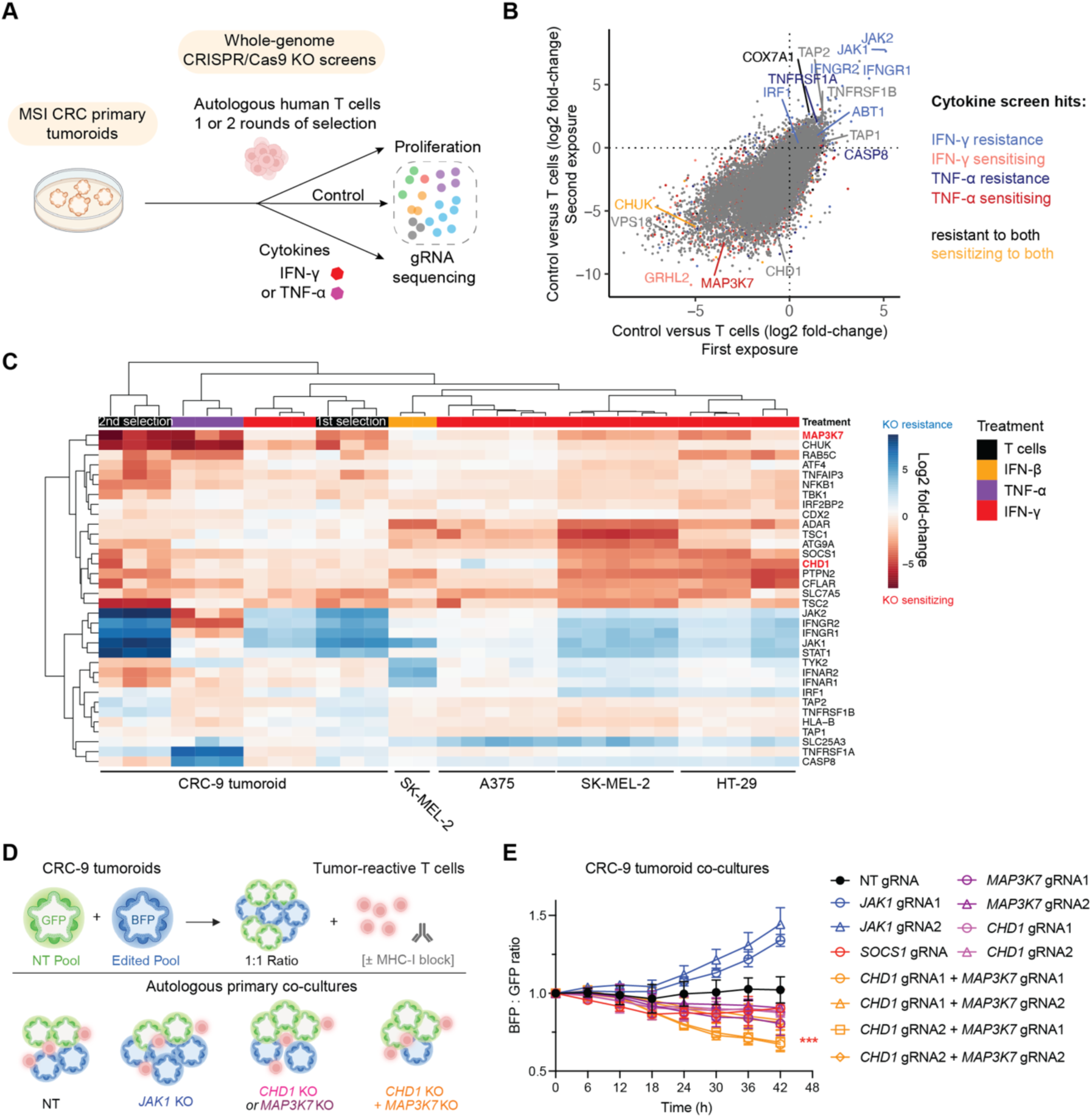
Autologous tumoroid-T cell co-culture CRISPR screens identify modulators of sensitivity to tumor-reactive T cells. **A)** Overview of the autologous co-culture CRISPR screens with primary tumoroids (CRC-9) and anti-tumor T cells. Tumoroids screens were performed ± IFN-γ (200 ng/mL) or TNF-*⍺* (100 ng/mL), or in the presence of tumor-reactive T cells (1:1 Effector: Target ratio) for 10 days. Tumoroids had one or two rounds of selection with T cells. MSI CRC; microsatellite-unstable colorectal cancer. **B)** Genetic modulators of cancer cell sensitivity to autologous human tumor-reactive T cells. Scatter plot comparing CRISPR KO screen log2-fold change (control versus T cells) from the first and second round of T cell selection. Data are representative of two independent screens performed on separate days. *CHD1* and selected co-culture hits from cytokine tumoroid screens are highlighted (*P* adjusted < 0.05). Pearson correlation *r* = 0.68. **C)** Heatmap displaying clustering of cell models and immunological selection pressures based on CRISPR KO screen log2 fold-changes. Columns represent different CRISPR screens against the control sample (e.g. control versus interferon or WT versus *JAK1* KO in the presence of interferon). See also Figure S4B. **D)** Overview of a competition assay using patient-derived CRC tumoroid co-cultured with autologous tumor-reactive T cells. The ratio of CRISPR/Cas9 edited (BFP^+^ and mCherry^+^) and non-targeting gRNA harboring (GFP^+^ and mCherry^+^) tumoroids was monitored over time. **E)** *CHD1* and *MAP3K7* loss additively sensitize cancer cells to killing by autologous T cells. Fluorescence of the different cell populations in the competition assay were measured using an Incucyte. Two-way analysis of variance (ANOVA); \*\*\**P* <0.001.

Integrating tumoroid-T cell co-culture cytokine screens (Supplemental Note 1 and Fig. S4A) revealed that sensitivity to tumor-reactive T cells was highly dependent on IFN-γ signaling (*JAK1, JAK2, IFNGR1, IFNGR2, STAT1*), with the exception the TNF-*⍺* receptor (*TNFRSF1A*, *TNFRSF1B*; Fig. 2B). We also identified cytokine-independent hits in T cell screens, such as *TAP1/2* KO associated with resistance; genes essential for antigen processing and presentation^76^ (Fig. 2B-C and Fig. S4B-C). In addition, KO of a proposed endogenous neoantigen in this model, mutant *ABT1* (but not *EEF1A1*^71^), conferred resistance to T cells in one of two T cell batches. Notably, *CHD1* KO was not significantly depleted with T cell treatment, but *MAP3K7* KO sensitized CRC-9 tumoroids to anti-tumor T cells to a similar or greater extent to known regulators of anti-tumor immunity, such as *SOCS1*^12^, *ADAR*^77^, *PTPN2*^10^ and *TBK1*^78^ (Fig. 2B-C). *MAP3K7* was in the top 1 % of sensitizing hits in TNF-*⍺* screens, implying TNF-*⍺* dependency in this model. *CHUK* encodes IKK*⍺*, part of the IKK complex with MAP3K7 (TAK1) that regulates NF-*κ*B activity^34^. *CHUK* KO also sensitized to T cells, TNF-*⍺* and IFN-γ, emphasizing the importance of NF-*κ*B signaling in tumor cell susceptibility to T cells (Fig. 2C). In co-culture cell competition (Fig. 2D), both *CHD1* and *MAP3K7* KO sensitized to T cell-mediated killing to a similar degree to *SOCS1* KO (Fig. 2E). Sensitization was enhanced in dKO tumoroids, perhaps explaining why *CHD1* KO alone was not a significant hit in genome-wide T cell co-culture screens. Furthermore, dKO tumoroids were more sensitive to IFN-γ than WT controls (Fig. S4D). Collectively, these data suggest that *CHD1* and *MAP3K7* loss additively enhances cancer cell sensitivity to tumor-reactive T cells.

### *CHD1* and *MAP3K7* control the transcriptional response to IFN-γ

To investigate the mechanism through which *CHD1* and *MAP3K7* loss sensitizes to IFN-γ and anti-tumor T cells, we generated single and dKO HT-29 and VCaP cancer cell models (Fig. S3C and S3F) and performed RNA sequencing in the presence and absence of IFN-γ (Fig. S5A). This analysis confirmed decreased expression of *CHD1* and *MAP3K7* in KO samples, implying nonsense-mediated decay (Fig. 3A), and revealed the broad transcriptional impact of *CHD1* KO, reflecting its role in chromatin remodelling^30^. In HT-29, *CHD1* KO increased the expression of *IRF8*, a master regulator of IFN-γ signaling^79^, and *TNFRSF1B*, which encodes part of the TNF-*⍺* receptor. *MAP3K7* KO reduced *TNFAIP3* and *CXCL10* expression in both cell models, suggesting a degree of mechanistic overlap between tissue types and consistent with a conserved role for *MAP3K7* in NF-*κ*B signaling^34^. Pathway analysis^80,81^ revealed enrichment in genes relating to JAK-STAT signaling in single KO and dKO VCaP cells (Fig. 3B), although this was not the case in HT-29 cells, which displayed varying levels of decreased JAK-STAT signaling (Fig. S5B). TNF-*⍺* signaling via NF-*κ*B was the most downregulated pathway from gene set enrichment analysis (GSEA) of HT-29 dKO cells treated with IFN-γ for 72 h, but this was significantly upregulated in VCaP cells, reflecting tissue-specific effects (Table S2). However, both cell models displayed a decrease in NF-*κ*B signaling mediated through *MAP3K7* KO (Fig. 3B and Fig. S5B). Androgen receptor signaling was significantly upregulated in *MAP3K7* KO and *CHD1* KO conditions in VCaP (Fig. 3B), implying a convergent mechanism and consistent with the frequent co-deletion of these genes in prostate cancer^21,24^ (Fig. 3B and Supplemental Note 2).

**Figure 3.**
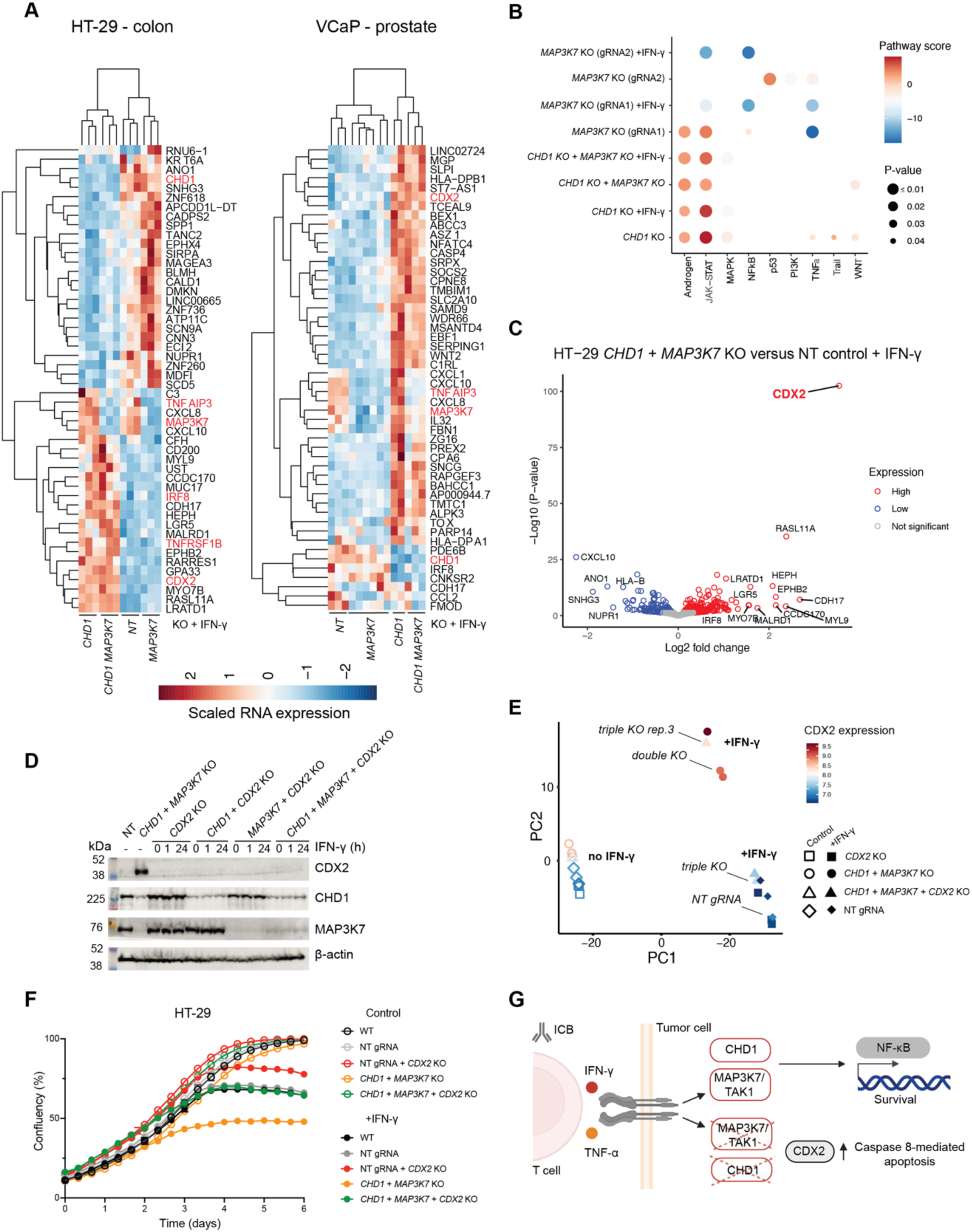
CHD1 and MAP3K7 control the response to IFN-γ in cancer cells through a transcriptional network dependent on CDX2. **A)** RNA sequencing analysis reveals transcriptional programs dependent on CHD1 and MAP3K7 in response to IFN**-**γ. Heatmap and hierarchical clustering of gene expression changes in *CHD1*, *MAP3K7* and dKO HT-29 and VCaP cells. Relative gene expression values are scaled normalized read counts from DESeq2. Columns represent independent biological replicates, with three biological replicates for each genotype except for VCaP *CHD1* KO (two replicates, Methods) and VCaP *MAP3K7* KO (six replicates, three replicates for two gRNAs). Differentially expressed transcripts of interest are highlighted in red for the top 50 significant transcripts (*P*-adjusted value < 0.05). **B)** Pathway analysis reveals altered NF-*κ*B, JAK-STAT and androgen receptor signaling in *CHD1* and *MAP3K7* KO VCap cells. Pathway activity for single and dKO cells ± IFN**-**γ (500 U/mL) for 24 h. *P* values and pathway scores are derived from decoupleR-PROGENy. **C)** *CDX2* expression is upregulated in dKO HT-29 cells. Volcano plot of differentially expressed genes comparing dKO and NT gRNA harboring HT-29 cells cultured in IFN-γ (500 U/mL) for 24 h. Data represent the average of three independent biological replicates. **D)** Confirmation of *CHD1* and *MAP3K7* KO and upregulation of *CDX2* in dKO cells. Western blotting of HT-29 cells with the indicated KO ± IFN-γ (500 U/mL) for 1 or 24 h. **E)** *CDX2* KO reverses IFN-γ-induced transcriptional programs in dKO cells. Principal component analysis comparing normalized RNA counts from *CDX2* KO, *CHD1* and *MAP3K7* dKO, triple KO (*CDX2* + *CHD1* + *MAP3K7* KO), and NT gRNA harboring HT-29 cells grown ± IFN-γ (500 U/mL) for 72 h before analysis. **F)** KO of *CDX2* rescues the IFN-γ sensitization effect of *CHD1* and *MAP3K7* KO in HT-29 cells. Cell proliferation was monitored ± IFN-γ (500 U/mL) using an Incucyte. Data represent the mean ± S.D of three technical replicates and are representative of two independent experiments. **G)** Model outlining how tumor cell *CHD1* and *MAP3K7* loss coordinately sensitizes to T cells and lymphocyte-derived cytokines. *CHD1* and *MAP3K7* loss alters cancer cell transcriptional response to cytokines through upregulation of the transcription factor CDX2 and reduced NF-*κ*B signaling.

### CDX2 mediates IFN-γ sensitivity induced by *CHD1* knockout

The most significant gene expression change in IFN-γ-stimulated dKO HT-29 cells compared to NT gRNA control cells was induction of *CDX2* (Fig. 3C), a transcription factor involved in NF-*κ*B signaling, tissue inflammation and development^82^. Notably, CDX2-positive CRC has a better prognosis^83^ and is associated with higher ICB response rates^84^ and *CDX2* KO caused resistance to IFN-γ in CRC-9 tumoroid screens (Fig. S4B). Increased *CDX2* expression was predominantly driven by *CHD1* KO with some contribution from *MAP3K7* KO (Fig. S3D) and was evident in both cell models but more pronounced in HT-29. This could reflect tissue-specific differences in baseline *CDX2* expression, which were higher in VCaP cells (Fig. S3D and Fig. S5C-D). Western blot analysis verified the induction of CDX2 in *CHD1* and *MAP3K7* KO HT-29 and VCaP cells at the protein level (Fig. 3D, Fig. S3D and Fig. S5D).

To investigate the potential role of CDX2 in regulating IFN-γ response, we generated a triple KO (tKO) HT-29 cell model deficient in *CDX2* (Fig. 3D). RNA-seq analysis of *CDX2* KO cells revealed significant downregulation of *IRF8* (Fig. S5E) and the IFN-γ pathway in GSEA (Fig. S5F). Principal component analysis of RNA expression revealed clustering by genotype, with tKO cells clustering with NT gRNA control samples, implying partial reversion of the induced transcriptional signature in dKO cells (Fig. 3E). One tKO biological replicate (rep.3) had higher levels of *CDX2* (Fig. S5G) and clustered with dKO samples (Fig. 3E), suggesting a threshold level of *CDX2* expression is required to sustain this transcriptional program.

Transcription factor analysis highlighted increased CDX2 activity in dKO cells, reduced NF-*κ*B activity (REL, RELA, NFKB1) and decreased activity of transcription factors involved in vesicular trafficking, autophagy (ATF2, ATF4) and IRF/STAT (Fig. S6A). Consistently, GSEA revealed increased mTORC pathway activity and decreased autophagy - a known immune evasion mechanism^13^, which was reversed with *CDX2* KO (Fig. S6B). Moreover, *CDX2* KO reversed gene programs induced in the dKO genotype, including TNF-*⍺* signaling via NF-*κ*B (Fig. S6C-D). *CDX2* KO was protective against IFN-γ in VCaP cells but could not fully reverse IFN-γ sensitization in the dKO context (Fig. S6E). In contrast, *CDX2* KO fully rescued sensitization to IFN-γ in HT-29 (Fig. 3F). tKO cells had comparable sensitivity to IFN-γ as NT gRNA control cells, consistent with *CDX2* playing a key role in sensitizing HT-29 cells to IFN-γ (Fig. 3G).

### *Chd1* and *Map3k7* deletion enhances anti-tumor immunity and response to immune checkpoint blockade in a mouse model

To assess the potential relevance of our findings *in vivo*, we selected a mouse model of melanoma, B16-F10, due to its immunogenicity and well-validated response to immunotherapy^85^. We generated a B16-F10 model deficient in *Chd1* and *Map3k7* (Fig. S7A) and subcutaneously engrafted control cells (expressing a NT gRNA) or dKO cells into syngeneic C57BL/6 mice. Although dKO cells grew at a comparable rate to control cells *in vitro* (Fig. S7B), engraftment rates were lower (Fig. 4A) consistent with a non-cell autonomous anti-tumor effect. Furthermore, dKO cells grew slower *in vivo* compared to controls (Fig. 4B).

**Figure 4.**
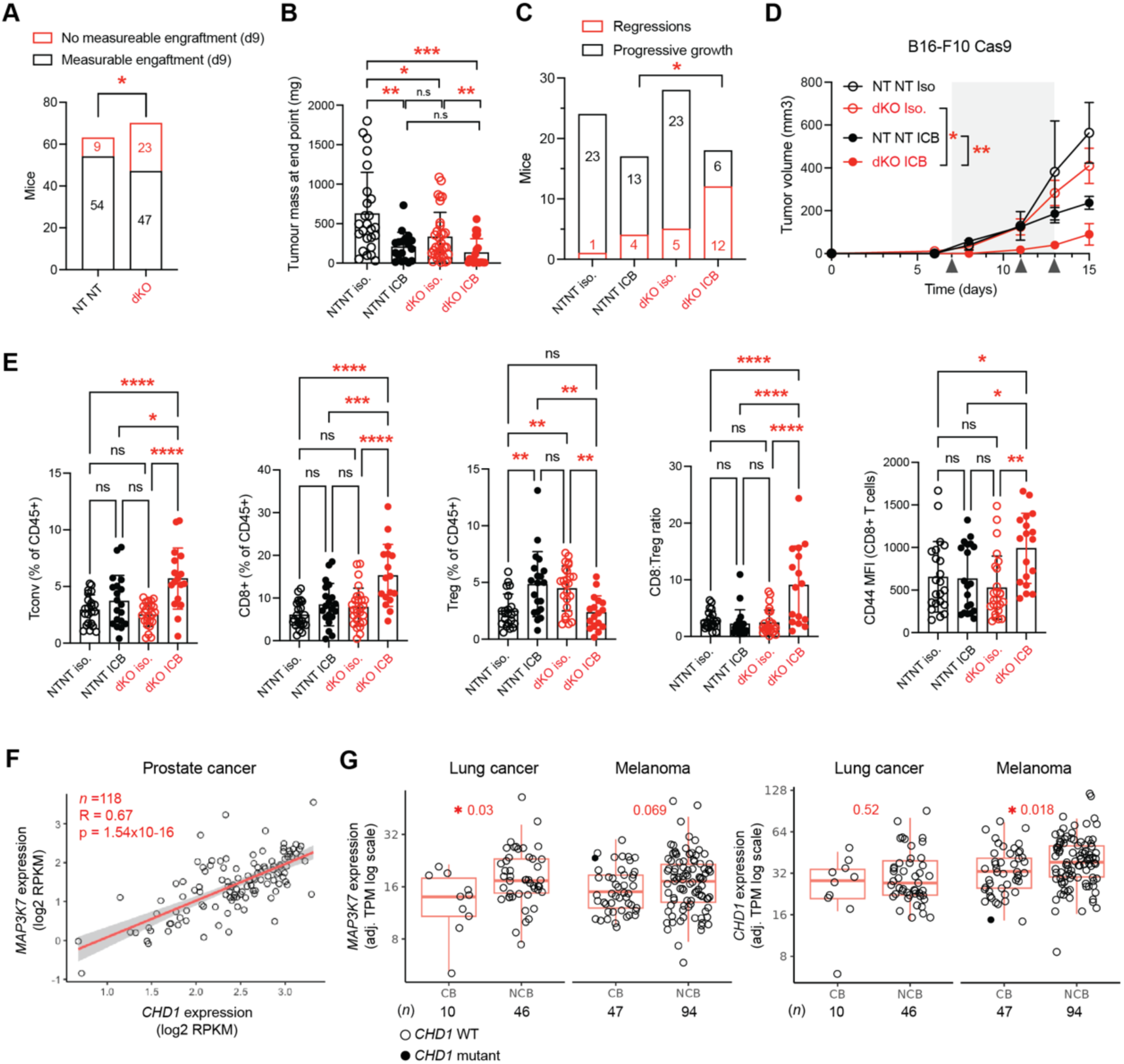
Reduced *CHD1* and *MAP3K7* expression enhances anti-tumor immunity and correlates with clinical response to immune checkpoint blockade. **A)** *Chd1* and *Map3k7* dKO B16-F10 tumor cells have higher frequency of spontaneous rejection. Engraftment rates of NT gRNA harboring or dKO B16-F10 cells injected subcutaneously into syngeneic C57BL/6 mice at day nine post-injection. Data represents four independent experiments. Two-sided Fisher’s exact test; * *P* = 0.015. **B)** *Chd1* and *Map3k7* dKO B16-F10 tumors grow slower *in vivo*. Endpoint tumor mass of successfully engrafted B16-F10 NT gRNA and dKO tumors in mice treated with intraperitoneal anti-PD-1 and anti-CTLA-4 immune checkpoint blockade (ICB) or isotype (Iso.) control. Data represent the mean ± S.D from three independent experiments. Unpaired, two-tailed Student’s t-test; *n*=25 NT NT iso.; *n*=18 NT NT ICB; *n*=33 dKO iso.; *n*=21 dKO ICB. \*\*\**P* = 0.0002, \*\**P* = 0.0025 NT NT iso. versus NT NT ICB,\*\**P* = 0.0094 dKO iso versus dKO ICB, \**P* = 0.01. **C)** Tumor regressions are more frequent in *Chd1* and *Map3k7* dKO B16-F10 tumors treated with ICB. Subcutaneous tumors were measured using calipers and tumors with a decrease in volume were considered as a regression. Data are pooled from three independent experiments. NT NT ICB versus dKO ICB, two-sided Fisher’s exact test; * *P* = 0.0176. **D)** Improved immunotherapy response in *Chd1* and *Map3k7* dKO B16-F10 tumors. Growth curves of subcutaneously engrafted B16-F10 NT NT gRNA or dKO tumors ± ICB. Data represent the mean ± S.E.M and are representative of three independent experiments. Two-way analysis of variance (ANOVA); \**P* = 0.0094; \*\**P* = 0.0019. dKO Iso *n*=6, dKO ICB *n*=5, NT NT Iso *n*=4, NT NT ICB *n*=6. **E)** Heightened anti-tumor immunity in *Chd1* and *Map3k7* dKO B16-F10 tumors treated with ICB. Immunoprofiling of subcutaneous tumors at the endpoint (day 15-17). Relative abundance of intratumoral T conventional CD4^+^, CD8^+^, T regulatory (T_regs_) and the CD8^+^:T_reg_ ratio, and surface expression of the activation marker CD44 on CD8^+^ T cells was assessed by flow cytometry. MFI; mean fluorescence intensity. Data represent the mean ± S.D and are pooled from three independent experiments. One-way ANOVA; \*\*\*\**P* < 0.0001; \*\*\**P* <0.0005; \*\**P* <0.01; \**P* <0.05; ns = not significant. NT NT Iso. *n*=22 or *n*=23 for CD8 analysis. NT NT ICB *n*=20 or *n*=19 for Tconv. analysis. dKO Iso. *n*=25 or *n*=27 for CD8 analysis. dKO ICB *n*=17 or *n*=16 for CD8:Treg or *n*=18 for CD44 analysis. **F)** Correlation of *CHD1* and *MAP3K7* mRNA expression in prostate cancers^86^ expressed as log2 fragments per kilobase of transcript per million (FMPK). Spearman’s rank correlation, R = 0.67, *P* = 1.54e-16. **G)** Reduced expression of *CHD1* and *MAP3K7* in lung cancer and melanoma is associated with clinical response to ICB. Box-plot displaying mRNA expression of tumor *CHD1* and *MAP3K7* expression (adjusted transcripts per million; adj. TPM) and clinical responses to ICB in patients from the Hartwig Medical Foundation^87^. Significance was assessed using the Wilcoxon signed-rank test and *n* denotes the number of patients. Boxplots represent the median, interquartile range (IQR) and whiskers are the lowest and highest values within 1.5 × IQR. Clinical benefit, CB; no clinical benefit, NCB.

To test whether loss of tumor cell *Chd1* and *Map3k7* would alter responses to ICB, we treated tumor-bearing mice with a combination of anti-PD-1 and anti-CTLA-4 monoclonal antibodies to mimic ICB therapy regimes in melanoma patients^38^. We verified that systemic ICB was functional by measuring dendritic cell influx into inguinal, tumor-draining lymph nodes with flow cytometry (Fig. S7C-F). Both dKO and control tumors responded to ICB, however, dKO tumors regressed more frequently (Fig. 4C) and grew slower than control tumors on treatment (Fig. 4D). This was associated with an increase in intratumoral CD8^+^ T cells, conventional CD4^+^ T cells and a reduction in CD4^+^Foxp3^+^ regulatory T cells (T_regs_) in dKO tumors compared with control tumors treated with ICB, resulting in an elevated CD8^+^:T_reg_ ratio (Fig. 4E). Moreover, ICB-treated dKO tumors displayed an increase in activated (CD44^+^) intratumoral CD8^+^ T cells, overall indicating a heightened anti-tumor adaptive immune response following ICB (Fig. 4E and Supplemental Note 3).

### Reduced *CHD1* and *MAP3K7* expression correlates with response to immune checkpoint blockade in patients

To evaluate the potential clinical relevance of our findings, we compared tumor *CHD1* and *MAP3K7* mRNA expression levels in patient samples. In prostate tumors, where dysregulation of these genes is most prevalent^22^, *CHD1* and *MAP3K7* mRNA expression were highly correlated (R = 0.67), reinforcing the concept of co-regulation^86^ (Fig. 4F). Interestingly, *JAK2* was significantly co-expressed with *CHD1* and *MAP3K7* (Fig. S8A), implying coregulation with key IFN-γ pathway genes. In patient records from the Hartwig Medical Foundation^87^ (HMF), *CHD1* and *MAP3K7* mRNA expression in tumors was significantly correlated with clinical responses to ICB, including anti-PD1/PD-L1 and anti-CTLA-4 (Fig. 4G, Methods). Lung cancer patients with clinical benefit from ICB had lower tumor *MAP3K7* expression (*P* = 0.03), and melanomas with clinical benefit from ICB had lower tumor *CHD1* expression than unresponsive patients (*P* = 0.018), outperforming known biomarkers of response such as *CD274* (PD-L1) and *CD8A* in lung cancer and *CCND1* ^88^ in lung cancer and melanoma (Fig. S8B-E and Supplemental Note 5). Taken together these data suggest that reduced tumor *CHD1* and *MAP3K7* expression could serve as biomarkers of ICB response.

## Discussion

Here we present a functional genomics landscape of genetic dependencies in four cancer cell models in the context of diverse immunological selection pressures, comprising 155 CRISPR screening samples. To map genetic determinants of sensitivity to T cells, we developed a co-culture CRISPR screening platform using autologous, tumor-reactive T cells and primary tumoroids expressing endogenous tumor neoantigens. Although our *in vitro* approach does not fully recapitulate the cell-cell interactions that occur *in vivo*^89^, our co-culture screening platform is more scalable than *in vivo* screens^90^, and by using primary human material, accounts for potential cross-species differences^91^.

Integrated analysis of CRISPR/Cas9 KO screens across cancer cell models and different cytokines facilitated deconvolution of contributions of each cytokine to tumor cell killing and identified shared and private genetic modulators of response, including a key role for mTOR signaling (Supplemental Note 4). We identified an acquired vulnerability in tumor cells that have lost *CHD1* and *MAP3K7* expression - enhanced sensitivity to IFN-γ, TNF-*⍺* and tumor-reactive T cells. *CHD1* and *MAP3K7* control cancer cell transcriptional responses to IFN-γ, at least in part, through regulating the expression of the transcription factor CDX2, and RNA-seq data indicates differing levels of *CDX2* expression might result in tissue-specific differences. *CHD1* and *MAP3K7* deletion has been shown to impact interferon response gene expression and sensitivity to oncolytic viruses in prostate cancer cells^92^. MAP3K7 and CHD1 are involved in activating NF-*κ*B signaling^93^ and transcription^32^ and CDX2 is itself regulated by NF-*κ*B^82^, collectively supporting a mechanism whereby *CHD1* and *MAP3K7* loss reduces NF-*κ*B signaling in response to cytokines, thereby priming cancer cells for apoptosis^94,95^ (Fig. 3G).

In summary, the functional genomics dataset presented here provides a rich resource for investigating the networks underlying cytokine signaling in inflammatory diseases and cancer immunity and highlights *CHD1* and *MAP3K7* loss as a potential biomarker of ICB response, thereby presenting future opportunities to improve cancer immunotherapy outcomes.

## Methods

This research was conducted in accordance with institutional guidelines at the Wellcome Sanger Institute and Cancer Research UK Cambridge Institute as outlined in the Good Research Practice Guidelines (v4, 2021) and Home Office project license number PP7993249. We support inclusive, diverse, and equitable research.

### Cancer cell line models

All cell line models used in this study (HT-29, A375, SK-MEL-2, VCaP) were verified as mycoplasma-free and STR profiled in accordance with authentication guidelines. Cells were cultured in RPMI or DMEM medium with 10% FCS and 1X penicillin-streptomycin or RPMI medium with 10% FCS, 2.5g Glucose, 1X Sodium Pyruvate and 1X penicillin-streptomycin (Thermo Fisher Scientific) at 37°C in a humidified incubator with 95% air and 5% CO_2_.

### *JAK1* and *JAK2* KO clone generation

Cas9 expressing cell lines HT-29, A375 and SK-MEL-2 were plated in a 6 well plate at a density to achieve 80% confluency after 24 h. After 24 h, transient transfection of *JAK1* or *JAK2* gRNAs was achieved by replacing media with 1.9 mL culture media, then combining 100 µL/well of Opti-MEM (Thermo Fisher Scientific), 1 ug/well of plasmid (gRNA-GFP, Addgene #48140), and 3 uL/well FuGENE (Promega) which was incubated at room temperature for 15 mins then added dropwise into the wells. After 24h incubation, media was refreshed, and cells incubated for a further 24 h. Cells were then stained with 1 µg/mL DAPI (Sigma-Aldrich) before sorting (DAPI^−^/GFP^+^) 1 cell per well into a 96 well plate containing their respective advanced media (advanced-RPMI and advanced-DMEM; Thermo Fisher Scientific). Clones were expanded and assessed for genomic *JAK1* or *JAK2* KO by PCR of the gRNA target site and TIDE analysis^96^. All primers are listed in Table S3. After 9 days, successful clones were expanded and assessed for GFP expression using Incucyte S3 (Sartorius) to ensure a lack of plasmid integration. GFP-negative clones were expanded further and frozen. Protein level JAK1/2 KO and Cas9 expression and abolition JAK-STAT signaling was assessed using western blot for JAK1/2, pSTAT1/3, and Cas9 as described later in the methods. Finally, functional resistance to the IFN-γ pathway was assessed by treating clones with 400 U/mL IFN-γ (Thermo Fisher Scientific - 300-02-100ug) and assessing their change in growth dynamics compared to wild type over 5-7 days using Incucyte S3 (Sartorius) confluency assessment. Analysis was performed using Incucyte S3 software (v2018B), phase confluency was determined using classic confluence segmentation.

### Guide RNA library

We used the Human MinLibCas9 library (Addgene #164896), which contains 37,722 guides targeting 18,761 protein coding genes (two guides per gene), with a further 200 non-targeting guides, with no GRCh38 perfect alignment and at most three 2nt-mismatch alignments, added to allow future benchmarking. The vector backbone for the library was modified from pKLV2-U6gRNA(BbsI)-PGKpuro2ABFP-W by cloning in the ccdB resistance gene cassette from pKLV1-fl-U6gRNA(BbsI)-ccdB-PGKpuro2ABFP to generate a modified pKLV2-U6gRNA(BbsI)-ccdB-PGKpuro2ABFP-W vector (Addgene #153033).

The library was delivered into electrocompetent cells (Endura, Lucigen) by electroporation with multiple parallel transformations to maintain library representation, before propagation in LB supplemented with 100 µg/mL ampicillin, shaking at 30°C overnight. For virus packaging, HEK293T cells were co-transfected with the library plasmid pool, psPAX2 (Addgene #12260) and pMD2.G (Addgene #12259) plasmids using FuGene HD (Promega) in Opti-MEM (Thermo Fisher Scientific). Viral particles were collected in media supernatant 72 h post-transfection, filtered and frozen.

### Whole-genome CRISPR KO screens and library preparation

To generate Cas9 expressing cell lines, cells were transduced overnight with lentivirus containing Cas9 (pKLV2-EF1a-Cas9Bsd-W; Addgene #68343) plus polybrene (8 μg/mL; Thermo Fisher Scientific). 24 h post-transduction, lentivirus-containing medium was refreshed with complete medium. 48 h post-transduction, positively transduced cells were selected for with blasticidin (Thermo Fisher Scientific). Cas9 activity was determined as described previously^97^. Briefly, cells were transduced with Cas9 reporter virus (pKLV2-U6gRNA5 (gGFP)-PGKmCherry2AGFP-W; Addgene #67982), as described above. The number of BFP^+^ and GFP-mCherry double-positive cells were determined by flow cytometry on a BD LSR Fortessa instrument (BD Biosciences), and data were subsequently analyzed using FCSExpress to determine the percentage of mCherry^+^ cells.

For CRISPR KO screening, cells were transduced with the MinLib virus (Addgene #164896) as described above, to achieve an infection rate of approximately 30%, as measured by BFP fluorescence with flow cytometry on a BD LSR Fortessa instrument (BD Biosciences), and an approximate 300 X coverage (estimated cells/gRNA). Cells were selected with puromycin (Thermo Fisher Scientific) for four days before taking a time 0 cell pellet. Screens were split into cytokine treatment arms and a control arm then cultured for a further 8 days, with passaging or refreshing cytokine every 3-4 days. Cytokine concentrations were based on IC50 CellTiter-Glo (Progema) viability drug titration assays or Incucyte S3 confluency assessment for concentrations reducing growth rate by 50%. Growth rate was determined using Incucyte S3 software (v2018B) by phase confluence with classic confluence segmentation. Cytokine concentrations used in screens were as follows: 400 U/mL IFN-γ (Thermo Fisher Scientific - 300-02-100ug), 400 U/mL IFN-β (STEMCELL Technologies), 20 ng/mL IL-6 (Thermo Fisher Scientific). Each screen was independently repeated twice on separate weeks.

Genomic DNA was extracted from cell pellets (DNeasy Blood and Tissue Kit; Qiagen) and the gRNA cassette was PCR amplified (PCR1: 28 cycles, 0.3 µM each primer, gDNA 3 µg/reaction using KAPA HiFi HotStart Ready Mix; Roche) with 24 reactions in parallel to maintain the complexity of the library. Each PCR1 product was QC checked by running on a gel to confirm that the bands were of similar strengths. Replicates were then pooled (5 µL removed from each and combined) and column purified (QIAquick PCR Purification Kit; Qiagen) before quantification using a Qubit - High Sensitivity kit (dsDNA Quantification Assay Kit; Thermo Fisher Scientific) and diluted to 200 pg/µL in nuclease free water. PCR1 products were then indexed (PCR2: 8 cycles, 0.2 µM each primer, 1 ng PCR1, using KAPA HiFi HotStart Ready Mix; Roche) and purified using AMPure beads (40µL of beads to 50µL sample; Beckman Coulter). Each PCR2 product was verified for purity and size using 2100 Bioanalyzer using the Bioanalyzer High Sensitivity DNA Analysis kit (Agilent). Libraries were sequenced on the HiSeq2500 (Illumina) using 19 bp SE sequencing on Rapid Run mode with a custom primer (9385110-U6-Illumina-seq2). All primers and PCR programs used are listed in Table S3.

### Whole-genome CRISPR KO screen analysis

To analyze CRISPR screens we used MAGeCK^98^ to generate comparisons between control and cytokine-treated arms. We compared WT, *JAK1* KO, *JAK2* KO and the independent cell clones of each experiment separately. We pooled replicates from two independent screens to generate an average read-count for each gRNA, with the MinLibCas9 library having two independent high-activity gRNAs per gene. As part of the implementation of MAGeCK, gRNAs were grouped together to give gene-level log2 fold-change values and relevant statistics. Any gRNAs with a count of 0 in the control samples were filtered out of the downstream analysis. We set a *P*-value cut off of < 0.05 to assess significance and an effect size of log2 fold-change of < −0.5 or > 0.5 for sensitizing or resistance gene hits, respectively. For tumoroid screens, FDR < 0.2 was used as an additional criterion to increase stringency due to higher signal-noise. R code used for downstream analysis is available on GitHub (https://github.com/MatthewACoelho/Watterson_etal_CRISPR_analysis). For pathway enrichment of CRISPR/Cas9 hits in cancer cell models, we used g:Profiler^99^.

### B16-F10 syngeneic transplants

B16-F10 Cas9 cells were generated by lentiviral infection of a Cas9 expressing construct under blasticidin selection. Cell line identity was verified using STR testing and cells and were mycoplasma-free. We generated two lentiviral gRNA vectors targeting the mouse *Chd1* and *Map3k7* and verified KO and dKO through dual infection by western blotting following puromycin selection. gRNA 2 for *Map3k7* and *Chd1* were chosen for downstream i*n vivo* experiments. As a control cell line, we used B16-F10 cells dual infected with a non-targeting gRNA (NT). Primers used to generate gRNAs are listed in Table S3.

*In vivo* experiments were performed under project license number PP7993249 and in line with Cancer Research UK Cambridge Institute institutional guidelines. 1×10^6^ B16-F10 Cas9 cells were subcutaneously injected in 100 µl endotoxin-free PBS into the flank of 10-13 week-old female C57BL/6 mice (JAX labs). Experiments were blinded and tumors were monitored with calipers every 2-3 days by facility staff. Seven days post-engraftment, mice were treated with 0.2 mg each (i.e. 10 mg/kg for a 20 g mouse) of isotype-matched control antibodies (BP0089, BE02060, BioXCell), or anti-PD1 (RMP1-14) and anti-CTLA-4 (UC10-4F10-11) antibodies (BP0146, BE0032, BioXCell), dosing every 2-3 days for a total of three doses.

### Immunophenotyping with flow cytometry

For analysis of single cell preparations from tumor-draining lymph nodes (tdLNs) and non-draining lymph nodes (ndLNs), inguinal tdLNs were dissected and removed from the tumor if attached and harvested into 500 µL RPMI. Inguinal ndLNs contralateral to the tumor were harvested into 500 µL RPMI (without serum added). Lymph nodes were chopped finely using scissors and tissue was digested with collagenase I (563 U/ml) and DNase I (0.225 mg/ml) in 500 µL total volume of RPMI. Samples were incubated for 30 minutes at 37 ℃ with agitation, then resuspended to obtain a single cell suspension, and pelleted at 600 g for 5 minutes at room temperature. Lymph nodes were resuspended in PBS supplemented with 2 % heat-inactivated FBS (Gibco). 100 µL of each lymph node sample was plated per well for staining and analysis by flow cytometry.

For analysis of single cell preparations from B16-F10 subcutaneous tumors, tumors were dissected, weighed and harvested into 1 mL RPMI. We only harvested tumors from mice that were macroscopically visible by dissection at endpoint. Tumors were chopped finely using scissors and decanted into a 15 mL Falcon tube. 3 mL of tissue digestion buffer containing collagenase I (563 U/ml) and DNase I (0.225 mg/ml), made in RPMI was added per tumor sample. Samples were incubated for 45 minutes at 37℃ with agitation. Samples were then strained through a 70 µm filter to obtain a single cell suspension and centrifuged at 1500 rpm for 5 minutes at 4℃. Red blood cell lysis (Qiagen) was done for 3 minutes at room temperature, followed by centrifugation and resuspension in 1 mL of PBS supplemented with 2 % heat inactivated FBS. 100 µL of each tumor sample was plated per well for staining and analysis by flow cytometry.

For staining, samples were incubated with 50 µL of surface antibody stain master mix containing anti-mouse CD16/32 (Thermo Fisher) to block Fc receptors, for 25 minutes at 4 ℃. Samples were then washed with 100 µL of PBS, centrifuged at 1500 rpm for 5 minutes at 4 ℃. Samples were fixed for 20 minutes at room temperature, washed to remove the fixative and permeabilized using the Foxp3/Transcription Factor Kit (Thermo Fisher). Samples were incubated with intracellular antibodies diluted in the permeabilization buffer and incubated overnight at 4 ℃. Samples were then washed with centrifugation at 1500 rpm for 5 minutes at 4 ℃ and resuspended in 300 µL of PBS containing 123 count eBeads (Thermo Fisher) to determine absolute cell number.

Surface antibodies used for identification of lymphoid cell populations were CD45 BV510 (30-F11, Biolegend), CD3e PeCy7 (145-2C11, Thermo Fisher), B220 APC-eFluor 780 (RA3-6B2, Thermo Fisher), CD4 AF700 (GK1.5, eBioscience), CD8a BB700 (53-6.7, BD Biosciences), NK1.1 BUV 395 (PK136, BD Biosciences). Antibodies used to remove contaminating cell types for each flow cytometry panel were labelled “lineage” on flow plots. For lymphoid phenotyping, antibodies were conjugated to eFluor 450 and were FceRIa (MAR-1, eBioscience), CD172α (P84, Biolegend), Siglec-F (1RNM44N, eBioscience), XCR1 (ZET, Biolegend), CD64 (X54-5/7.1, Biolegend), CD11b (M1/70, Biolegend), I-A/I-E (CI2G9, BD), CD11c (N418, eBioscience), F4/80 (BM8, eBioscience), Ly6G (1A8-Ly6g, eBioscience), Ly-6C (HK1.4, eBioscience). Lineage antibodies for myeloid phenotyping were conjugated to eFluor 450 and were CD3 (145-2C11), NK1.1 (PK136), CD5 (53-7.3), CD19 (1D3) and B220 (RA3-6B2), all on eFluor450 (eBioscience). Surface antibodies used for the identification of myeloid cell populations were FceRIa PerCP-eFluor710 (MAR-1, eBioscience), CD172α AF488 (P84, Biolegend), Siglec-F SB600 (1RNM44N, eBioscience), XCR1 BV650 (ZET, Biolegend), CD64 BV711 (X54-5/7.1, Biolegend), CD11b BV785 (M1/70, Biolegend), I-A/I-E BUV395 (CI2G9, BD), CD11c AF700 (N418, eBioscience), F4/80 APC-eFluor780 (BM8, eBioscience), Ly6G PE-eFluor610 (1A8-Ly6g, eBioscience), Ly-6C PE-Cy7 (HK1.4, eBioscience). Lineage antibodies for myeloid phenotyping were conjugated to eFluor 450 and were CD3 (145-2C11), NK1.1 (PK136), CD5 (53-7.3), CD19 (1D3) and B220 (RA3-6B2), all on eFluor450 (eBioscience). The viability dye used for lymphoid and myeloid panels was a fixable viability dye, UV 455 (eBioscience). Data was acquired on a BD Symphony instrument and analysed using FlowJo Version 10.10.0.

### Arrayed screen validation experiments

Individual gRNAs sequences were extracted from MinLib gRNAs (Addgene #164896), ordered as oligos (Sigma-Aldrich), and cloned using Golden Gate cloning. Our procedure made use of primers encoding a gRNA with BbsI overhangs and an additional G for hU6 RNApolIII transcription (Forward: 5ʹ-CACCGNNNNNNNNNNNNNNNNNNN-3ʹ and Reverse: 5ʹ-AAACNNNNNNNNNNNNNNNNNNC-3ʹ), annealed by boiling (100°C 5 mins) and slowly cooling to room temperature (0.1°C per second until reaching 25°C), and then ligated duplexes with a BbsI entry vector (Addgene #67974 or a hygromycin-GFP version) using BbsI-HF (NEB), T4 DNA ligase and buffer (NEB), 1X BSA (NEB) for 30X cutting (37°C for 5 mins) and ligating (16°C for 10 mins) cycles, before heat-shock transformation of Stable Competent *E*. *coli* (NEB - C3040I) and spreading on 100 µg/mL ampicillin agar plates overnight at 37°C. Colonies were picked and expanded in LB containing 100 µg/mL ampicillin before shaking at 30°C overnight. *E.coli* were then harvested and the plasmids were column purified (Qiagen) and sequences were verified via Sanger sequencing of the plasmid with a U6 promoter (Eurofins). All gRNAs are listed in Table S3. For virus packaging, HEK-293T cells were co-transfected with gRNA plasmid, psPAX2 and pMD2.G plasmids using FuGene HD (Promega) in Opti-MEM (Thermo Fisher Scientific). Viral particles were collected in culture supernatant 72 h post-transfection, filtered and frozen. Lines were transduced with the gRNA virus (plus 8 µg/mL polybrene; Thermo Fisher Scientific) and selected with antibiotics (puromycin or hygromycin), followed by western blotting to ensure gene knockout or knockdown as described later in the methods.

### Cell Titre-Glo and cell proliferation assays

Cells were plated in white opaque 96 well plates (Corning) in triplicate for each condition to be tested, allowed to attach for 24 h, and then treated with IFN-γ (Thermo Fisher Scientific - 300-02-100ug). Plates were imaged using Incucyte S3 (Sartorius) every 8 h and confluency was assessed using Incucyte S3 analysis software. Analysis was performed using Incucyte S3 software (v2018B), phase confluency was determined using classic confluence segmentation. In the case of VCaP lines, IFN-γ was refreshed halfway through the culture period given the length of the assay (8 days), all other lines were optimized for a 6-day assay. Additionally, given the morphology of the cell line, VCaP cells were unable to be optimized for confluency assessment. Following the end of the treatment period, cell viability was assessed using CellTiter-Glo assay (Promega). All media was removed and replaced with 50 μL base media to which 25 μL of CellTiter-Glo (Progema) was added. Plates were protected from light and incubated at room temperature for 30 minutes before measuring well fluorescence (Paradigm). CellTiter-Glo fluorescence values were normalized by calculating them as a percentage of the mean average of their control wells.

### Western blotting

Cells were lysed in sample loading buffer (8% SDS, 20% β-mercaptoethanol, 40% glycerol, 0.01% bromophenol blue, 0.2 M Tris-HCL pH 6.8 or NuPAGE LDS Sample Buffer 4x with 10% β-mercaptoethanol) and boiled (95°C) for 5 minutes before loading onto a NuPAGE 4-12% Bis-Tris gel (Thermo Fisher Scientific). Proteins were transferred to a PVDF membrane (Amersham) in Tris-Glycine Transfer Buffer (Sera Care) before blotting overnight with the following primary antibodies: β-tubulin (#T4026, Clone TUB2.1, 1:2,000 dilution) from Sigma-Aldrich, or β-actin (#4970, Clone 13E5, 1:1,000 dilution), CHD1 (#4351, Clone D8C2, 1:1,000 dilution), MAP3K7 (#4 5206, Clone D94D7, 1:500 dilution), STAT1 (#9172, 1:1,000 dilution), pSTAT1-Y710 (#7649, Clone D4A7, 1:1,000 dilution), JAK1 (#3344, Clone 6G4, 1:1,000 dilution), JAK2 (#3230, Clone D2E12, 1:1,000 dilution), Cas9 (#14697, Clone 7A9-3A3, 1:1,000 dilution), CDX2 (#3977, 1:1,000 dilution), vinculin (#13901, Clone E1E9V, 1:1,000 dilution), cleaved caspase 3 (#9661S, Clone Asp175, 1:1,000 dilution), and caspase 8 (#9746T, Clone 1C12, 1:1,000 dilution) from Cell Signaling. Secondary antibodies (anti-mouse and ant-rabbit) were conjugated to horseradish peroxidase (#NXA931V and #NA934V, Amersham, 1:2,000-1:5,000 dilution). Membranes were imaged using G-Box iChemi XL (Syngene) or ImageQuant 800 (Amersham) imagers. Ladder used in all blots was the ECL Rainbow Marker - Full range (Sigma-Aldrich).

### RNA-seq sequencing

Cells were plated to achieve an approximate 80-90% confluency at the point of harvest for RNA extraction. Plates were incubated for 1 day prior to starting the first treatment with IFN-γ at 500 U/mL (Thermo Fisher Scientific - 300-02-100ug). Cells were treated for 0, 1 and 24 h in one experimental set (HT-29 and VCaP CHD1-MAP3K7) and 0 and 72 h in the second experimental set (HT-29 CHD1-MAP3K7-CDX2). RNA was extracted (RNeasy, Qiagen), DNA removed with DNase I digest (Qiagen), and reverse transcription performed using poly dT priming (SuperScript IV, Thermo Fisher Scientific). Libraries were sequenced on the NovaSeq6000 (Illumina) using 150 bp SP sequencing. All RNA sequencing experiments were repeated independently two-three times on separate days and averaged as indicated in figure legends. All primers are listed in Table S3.

### RNA-sequencing analysis

Paired-end transcriptome reads were quality filtered and mapped to GRCh38 (Ensembl build 98) using STAR-v2.5.0c^100^ with a standard set of parameters (https://github.com/cancerit/cgpRna). Resulting bam files were processed to get per gene read count data using HTSeq 0.7.2, which was used for downstream analysis. Using the DESeq2 R library^101^, merged per gene read counts were filtered to remove duplicates and genes which had <20 counts in ≥6 samples before undergoing differential expression and principal component analysis (https://github.com/ABWatterson/DESeq2_All_RNAseq_PCA and https://github.com/MatthewACoelho/Watterson_etal_RNAseq_analysis). Differential expression analysis was conducted by comparing all samples to one another and plotted using -log10 p-value vs Log2 Fold Change. Normalized and log transformed counts generated in the same DESeq2 pipeline were then used for further analyses. Using the decoupleR R library^81^, pathway and transcription factor activities were determined. For pathway analysis (https://github.com/ABWatterson/decoupleR-Pathway-activation-main), pathway activity was determined using the top 500 genes per pathway in the PROGENy model. For transcription factor analysis (https://github.com/ABWatterson/decoupleR-TF-activation), transcription factor activity was determined using the CollecTRI network^102^. Gene Set Enrichment Analysis was performed using software from the Broad Institute (GSEA version 4.3.3^103,104^) using standard parameters with no collapse (https://www.gsea-msigdb.org/gsea/doc/GSEAUserGuideFrame.html) and normalized counts.

### Autologous tumoroid - T cell and co-culture and cytokine CRISPR screens

Similar to our previously described method^36^, tumoroids were dissociated into single cells and incubated overnight in suspension culture with complete media containing pKLV2-EF1a-BsdCas9-W lentiviral particles and polybrene (8 μg/ml) to express Cas9. The following day, the cells were seeded in BME and cultured as tumoroids. Blasticidin selection (20 mg/ml) was initiated 48 h post-transduction and continued for the duration of the experiment. The tumoroid demonstrated Cas9 activity exceeding 80%. gRNAs from the minimal genome-wide human CRISPR-Cas9 library (MinLibCas9) were utilized^36^. Tumor tumoroids dissociated into single cells, and a total of 3.3 × 10⁷ cells were transduced overnight in suspension with the lentiviral-packaged whole-genome gRNA library. The transduction was performed at 30% efficiency to ensure 200× library coverage, with polybrene (8 μg/ml) included. To ensure high cell yields, tumoroids were cultured in suspension with 5% extracellular matrix (ECM) as previously described^74^. After 48 hours, tumoroids underwent puromycin selection (2 mg/ml). Fourteen days later, approximately 2 × 10⁷ cells were collected, pelleted, and stored at −80°C for DNA extraction. Additional cell pellets were collected at key time points: prior to cytokine or T cell exposure (T0), after cytokine treatment (200 ng/mL IFN-γ or 100 ng/mL TNF-*⍺*), and following the first and second rounds of T cell-mediated killing. Library transduced tumoroids were then stimulated with IFN-γ or TNF-*⍺* or left unstimulated as a negative control for 9-10 days. For T cell screens, the killing assay was designed to achieve approximately 50% tumoroid elimination, enabling the identification of genes associated with resistance and sensitization. The effect of T cells was initially titrated in a smaller format and subsequently scaled up to reach the desired killing efficiency. Tumoroids underwent two rounds of T cell exposure, each conducted at an optimized target-to-effector ratio of 1:1. T cell selections (killing assays) were conducted across multiple 6-well plates and lasted for 72 hours. Depleting nicotinamide from the complete CRC tumoroid medium had only a mild impact on tumoroid growth over a 3-day period and significantly improved their viability over one week compared to culturing in T cell medium. Importantly, nicotinamide depletion did not compromise the T cell killing capacity of CRC-9 tumoroids. Based on these findings, all screens were conducted using an tumoroid medium depleted of nicotinamide to maintain tumoroid viability, while ensuring optimal T cell function. After each selection, cells were reseeded in 5% ECM-supplemented media and cultured until reaching a minimum of 4 × 10⁷ cells, enabling subsequent selections and pellet collection for analysis. Genomic DNA was extracted using the Qiagen Blood & Cell Culture DNA Maxi Kit (13362) following the manufacturer’s protocol. PCR amplification, Illumina sequencing (19-bp single-end sequencing with custom primers on the HiSeq2000 v.4 platform), and sgRNA quantification were performed as previously described.

### Autologous tumoroid - T cell co-culture validation experiments

CRC-9 tumor tumoroid co-culture assays were performed as described^73^. CRC-9 Cas9-mCherry cells with NT gRNA (mAzami Green) were 1:1 co-cultured with cells with KO gRNA (BFP) in a competition assay. Briefly, 96 and 24 well-plates were coated with anti-CD28 antibody (2.2 μg/mL), then PBMCs were thawed and cultured at 2e6 cells per well in the 24 well-plate with T cell thawing media (RPMI, 1% penicillin-streptomycin, 1% Glutamax, 10% FBS (Gibco), and 150 U/mL IL-2 (Thermo Fisher Scientific)). On the same day, autologous CRC-9 tumoroids were removed from their 5% basement membrane extract (BME) (R&D systems) and replated with 0% BME and IFN-γ (Thermo Fisher Scientific) at 6e5 cells per well in 2 mL culture media (RPMI, 1% penicillin-streptomycin, 1% Glutamax, (Gibco), and 10% Human Serum (Sigma-Aldrich)). The following day, tumoroids (5,000 cells of BFP gRNA and 5,000 cells of mAzami Green gRNA) and T cells were plated in suspension at a 3:1 effector:target ratio for 72 h. Plates were placed in an Incucyte S3 (Sartorius) and wells were imaged every 6 h with phase and fluorescent optics for mCherry and mAzami Green. All assays were performed in the presence of the anti-PD-1 antibody nivolumab (20 µg/mL; Selleckchem) in culture media supplemented with primocin (Invivogen). Relative T cell killing was quantified by comparing the relative cell numbers between the BFP-gRNA (mCherry only) and NT-mAzami Green-gRNA (mCherry and mAzami Green) to give ratios which were normalized to the 0 h measurements. Analysis was performed using Incucyte S3 software (v2018B), mAzami Green and mCherry cell numbers were determined using Top-Hat segmentation.

### Analysis of publicly available cancer patient data

Studies in a variety of cancer types were selected from clinically available data that met the criteria of having patients profiled by WGS or RNAseq. Clinical data was acquired from cBioPortal^105–107^ (TCGA, Other studies). Using cBioPortal, studies were queried for deep deletion status of *CHD1* and *MAP3K7* in all samples. Samples identified to have a deep deletion in one or both genes were counted and then normalized as a percentage of the total number of samples of that cancer type within the study. Criterion for study inclusion in the final figure was <1% deep deletion in either or both genes. Study data used in this analysis are listed in Table S4.

### Hartwig Medical Foundation data analysis

Patients receiving “Immunotherapy” as their “consolidatedTreatmentType” were selected from the isofox RNA sequencing dataset (464/3,682). Clinical benefit (CB) was determined as a “firstResponse” of “PR” (partial response), “CR” (complete response). No clinical benefit (NCB) was defined as a “firstResponse” of “SD” (stable disease), “PD” (progressive disease), or “Clinical progression”, in line with previous reports^88^. We grouped cohorts by primary tumor location and filtered on groups > 50 patients on ICB; melanoma/skin, urothelial tract and lung. For RNA analysis we used normalized TPM (adjusted transcripts per million) plotted on a log-scale. All data analysis was performed using virtual machines with R and Unix on Google Cloud Platform in accordance with the Data Access Agreement with HMF. The Purple pipeline was used to extract mutations in driver genes, which included *CHD1* but not *MAP3K7*. ICB was anti-PD-1 (*n* = 53), anti-PD-L1 (*n =* 2) or anti-PD-1/CTLA-4 combination (*n* = 1) for lung cancer, and anti-PD-1 (*n* = 100), anti-CTLA-4 (*n* = 6) or anti-PD-1/CTLA-4 combination (*n* = 35) for melanoma. 1/141 melanoma patients treated with ICB had a deletion of *CHD1* in their tumor and clinical response.

### Statistics and Reproducibility

For multiple paired t-test, Two-tailed unpaired Student’s t-test, two-sided Fisher’s exact test and Wilcoxon signed-rank test, significance was defined as *P* < 0.05. No statistical method was used to predetermine sample size, nor were the experiments randomized. The investigators were not blinded to allocation during experiments and outcome assessment. For MAGeCK analysis of CRISPR screens, gRNAs with 0 counts in control samples were filtered out of downstream analysis. For bulk RNA-seq analysis, exclusion criteria were RNAs with low read counts (genes which had <20 counts in ≥6 samples) or replicates with Pearson’s correlation <0.9 and poor clustering by Euclidean distance between sample replicates, meaning we excluded VCap *CHD1* gRNA2 IFN-γ replicate 3 (Fig. S6A).

## Supplemental Tables

Table S1. Processed data from CRISPR KO screens

Table S2: RNA sequencing GSEA analysis

Table S3: Primers and gRNA sequences

Table S4: TCGA study sample data

Table S5: DNA and RNA sequencing ENA accession numbers

## Data availability

Sequencing data are deposited on ENA and accessions are listed in Table S5.

## Code availability

Code used to analyze CRISPR/Cas9 KO screens can be found on GitHub here: https://github.com/MatthewACoelho/.

Code used to analyze RNAseq data for differential expressions and principal component analysis can be found on GitHub here: https://github.com/ABWatterson/DESeq2_All_RNAseq_PCA

Code used to analyze RNAseq data for PROGENy model based pathway activation can be found on GitHub here: CollecTRI network https://github.com/ABWatterson/decoupleR-Pathway-activation-main

Code used to analyze RNAseq data for CollecTRI network based transcription factor activation can be found on GitHub here: https://github.com/ABWatterson/decoupleR-TF-activation

## Inclusion and Ethics

We support inclusive, diverse, and equitable research.

## Acknowledgements

This research was funded in whole, or in part, by the Wellcome Trust Grant 206194. This work was funded by Open Targets (OTAR2061). The authors wish to acknowledge the contribution of the Cancer Ageing and Somatic Mutation Support team at the Wellcome Sanger Institute. Figure components were created with BioRender.com. This work was supported by Cancer Research UK [RCCCDF-Nov23/100002]. This work was partly funded by a Sanger Accelerator Award for Postdocs, which included a contribution from Sanger’s portion of the UKRI Talent & Research Stabilisation Fund. For the purpose of Open Access, the author has applied a CC BY public copyright license to any Author Accepted Manuscript version arising from this submission. We thank Olli Dufva and Saroor Patel for critical reading of the manuscript and the Cancer Genome Editing laboratory for feedback on the manuscript. This publication and the underlying study have been made possible partly based on data that Hartwig Medical Foundation and the Center of Personalised Cancer Treatment (CPCT) have made available to the study through the Hartwig Medical Database. The data for this study are provided (in part) by the Netherlands Cancer Institute (Antoni van Leeuwenhoek ziekenhuis). Additionally, patient results published here are in part based upon data generated by the following initiatives: Therapeutically Applicable Research to Generate Effective Treatments (TARGET) initiative, phs000218.v24.p8, managed by the NCI (available at [National Cancer Institute (NCI) TARGET: Therapeutically Applicable Research to Generate Effective Treatments](https://www.ncbi.nlm.nih.gov/projects/gap/cgi-bin/study.cgi?study_id=phs000218.v24.p8)), the TCGA Research Network (https://www.cancer.gov/tcga), as well as [The Metastatic Breast Cancer Project](https://www.mbcproject.org/) and The [Metastatic Prostate Cancer Project](https://mpcproject.org/) projects of [Count Me In](https://joincountmein.org/). The authors want to thank all the involved patients, medical staff and research staff for making this possible.

## Author Contributions

MAC, MJG, TH and EEV devised the study. AW performed CRISPR and RNA-seq experiments and assisted with RNA-seq analysis. GP and SC devised tumoroid experiments and GP, SFV and AW executed tumoroid experiments. EK, SB, AW and MAC analyzed CRISPR screening and RNA-seq data. VV and CC developed tumoroid and autologous T cell reagents and protocols. TB assisted with patient data analysis. YS assisted with in vivo experiments, flow cytometry and analysis and TH advised on in vivo experiments and analysis. MAC performed HMF and CRISPR analysis. Funding acquisition was by MAC, MJG and EEV. MAC and MJG wrote, and all authors reviewed the manuscript.

