## Supplemental Figures for "CRISPR screens in the context of immune selection identify *CHD1* and *MAP3K7* as mediators of cancer immunotherapy resistance"

Figures S1-8.

**Figure S1**

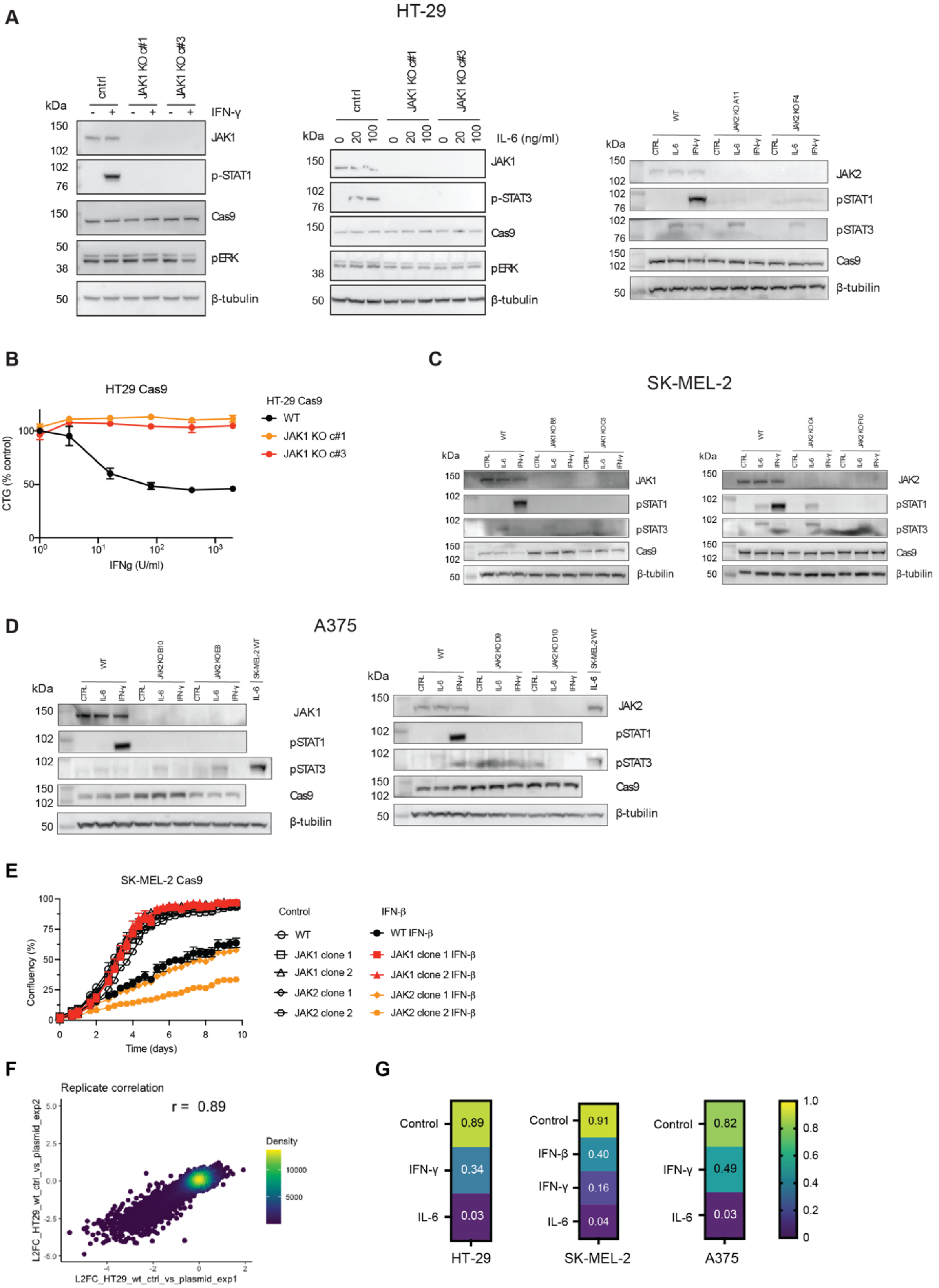
A
HT-29
kDa
cntrl
JAK1 KO c#1
JAK1 KO c#3
IFN-γ
JAK1
p-STAT1
Cas9
pERK
β-tubulin
IL-6 (ng/ml)
p-STAT3
JAK2
pSTAT1
pSTAT3
B
HT29 Cas9
CTG (% control)
IFNg (U/ml)
HT-29 Cas9
WT
JAK1 KO c#1
JAK1 KO c#3
C
SK-MEL-2
D
A375
E
SK-MEL-2 Cas9
Confluency (%)
Time (days)
Control
IFN-β
WT
JAK1 clone 1
JAK1 clone 2
JAK2 clone 1
JAK2 clone 2
WT IFN-β
JAK1 clone 1 IFN-β
JAK1 clone 2 IFN-β
JAK2 clone 1 IFN-β
JAK2 clone 2 IFN-β
F
Replicate correlation
r = 0.89
Density
L2FC_HT29_wt_ctrl_vs_plasmid_exp2
L2FC_HT29_wt_ctrl_vs_plasmid_exp1
G
Control
IFN-γ
IL-6
IFN-β
0.89
0.34
0.03
0.91
0.40
0.16
0.04
0.82
0.49
0.03
HT-29
SK-MEL-2
A375

**Figure S1.**

**A)** Confirmation of *JAK1* and *JAK2* KO in HT-29 *JAK1* and *JAK2* KO isogenic clone pairs. Western blotting of HT-29 cells in the presence or absence of IFN-γ (400 U/mL) or IL-6 (20 ng/mL) for 1 h before analysis.

**B)** *JAK1* KO renders HT-29 cells resistant to IFN-γ induced killing demonstrating functional KO. Cells were grown untreated or treated with IFN-γ (400 U/mL) for 5 days, after which viability was measured using Cell Titer-Glo (CTG). Data was normalized to untreated samples and represent the mean ± S.D. of three biological replicates.

**C)** Confirmation of *JAK1* and *JAK2* KO in SK-MEL-2 *JAK1* and *JAK2* KO isogenic clone pairs. Western blotting of SK-MEL-2 cells in the presence or absence of IL-6 (20 ng/mL) or IFN-γ (400 U/mL) for 1 h before analysis.

**D)** Confirmation of *JAK1* and *JAK2* KO in A375 *JAK1* and *JAK2* KO isogenic clone pairs. Western blotting of A375 cells in the presence or absence of IL-6 (20 ng/mL) or IFN-γ (400 U/mL) for 1 h before analysis.

**E)** Response of *JAK1* and *JAK2* KO SK-MEL-2 clones to IFN-β. Cell growth assays in the presence or absence of IFN-β (400 U/mL). Proliferation was monitored using an Incucyte. Data represent the mean ± S.D of three technical replicates.

**F)** Scatter plot of replicate correlations from CRISPR/Cas9 KO screens in HT-29. Log2 fold-change of the control samples versus the plasmid library is shown. Results were generated using MAGeCK and correlation determined using Pearson correlation coefficient (*r*).

**G)** Heatmap summary of replicate correlation for three cancer cell model CRISPR/Cas9 KO screens in the presence or absence of the indicated cytokine. Results were generated using MAGeCK and correlation determined using Pearson correlation coefficient (*r*) between independent screen replicates performed on different days.

### Figure S2

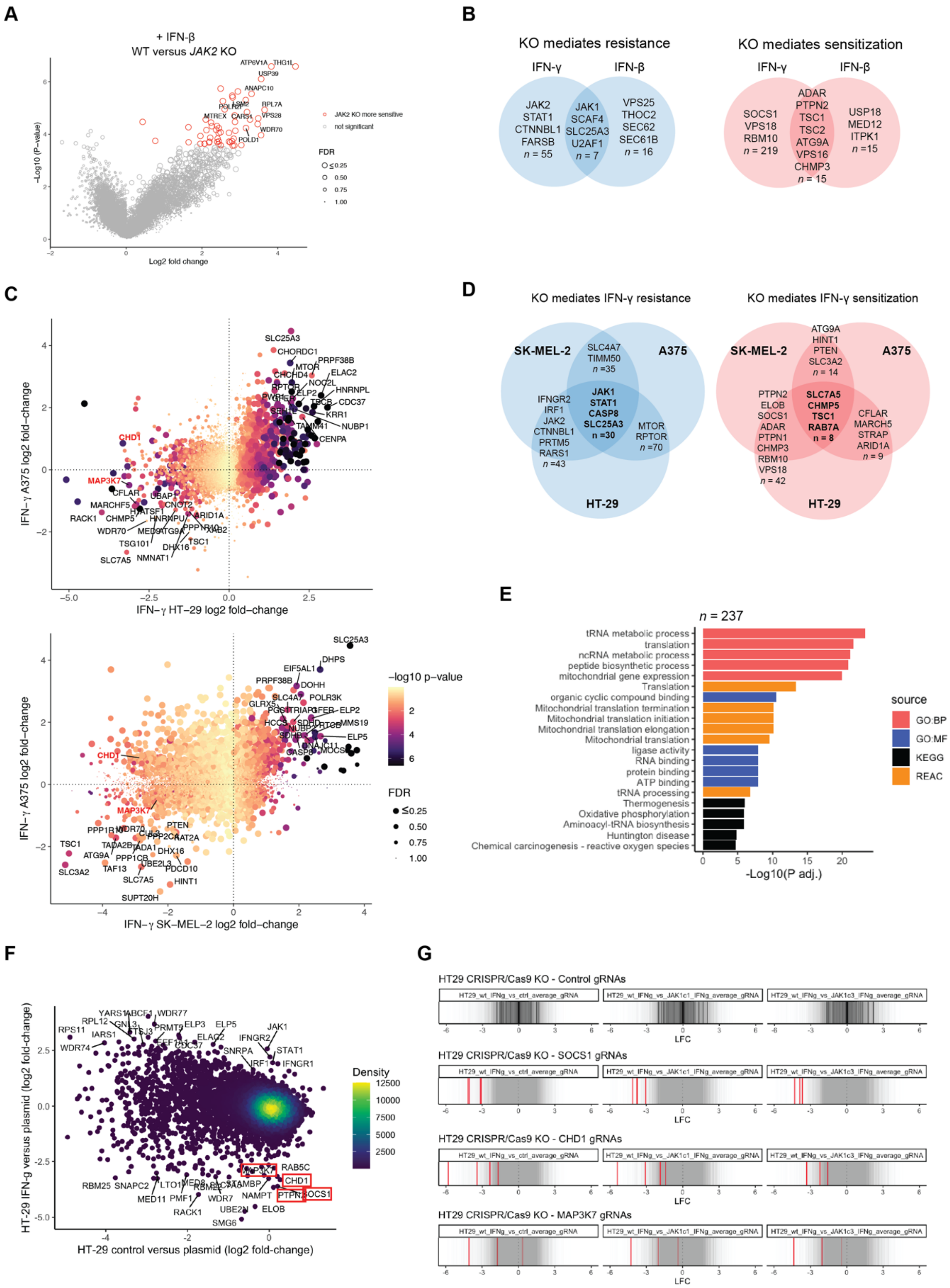

A
+ IFN-β
WT versus JAK2 KO
Log2 fold change
−Log10 (P-value)
JAK2 KO more sensitive
not significant
FDR
B
KO mediates resistance
IFN-γ
IFN-β
JAK2 STAT1 CTNNBL1 FARSB n = 55
JAK1 SCAF4 SLC25A3 U2AF1 n = 7
VPS25 THOC2 SEC62 SEC61B n = 16
KO mediates sensitization
SOCS1 VPS18 RBM10 n = 219
ADAR PTPN2 TSC1 TSC2 ATG9A VPS16 CHMP3 n = 15
USP18 MED12 ITPK1 n =15
C
IFN−γ HT−29 log2 fold−change
IFN−γ A375 log2 fold−change
IFN−γ SK−MEL−2 log2 fold−change
−log10 p−value
FDR
D
KO mediates IFN-γ resistance
SK-MEL-2
A375
HT-29
SLC4A7 TIMM50 n =35
JAK1 STAT1 CASP8 SLC25A3 n =30
IFNGR2 IRF1 JAK2 CTNNBL1 PRTM5 RARS1 n =43
MTOR RPTOR n =70
KO mediates IFN-γ sensitization
ATG9A HINT1 PTEN SLC3A2 n = 14
SLC7A5 CHMP5 TSC1 RAB7A n = 8
PTPN2 ELOB SOCS1 ADAR PTPN1 CHMP3 RBM10 VPS18 n = 42
CFLAR MARCH5 STRAP ARID1A n = 9
E
n = 237
tRNA metabolic process
translation
ncRNA metabolic process
peptide biosynthetic process
mitochondrial gene expression
Translation
organic cyclic compound binding
Mitochondrial translation termination
Mitochondrial translation initiation
Mitochondrial translation elongation
Mitochondrial translation
ligase activity
RNA binding
protein binding
ATP binding
tRNA processing
Thermogenesis
Oxidative phosphorylation
Aminoacyl-tRNA biosynthesis
Huntington disease
Chemical carcinogenesis - reactive oxygen species
-Log10(P adj.)
source
GO:BP
GO:MF
KEGG
REAC
F
HT-29 control versus plasmid (log2 fold-change)
HT-29 IFN-g versus plasmid (log2 fold-change)
Density
G
HT29 CRISPR/Cas9 KO - Control gRNAs
HT29 CRISPR/Cas9 KO - SOCS1 gRNAs
HT29 CRISPR/Cas9 KO - CHD1 gRNAs
HT29 CRISPR/Cas9 KO - MAP3K7 gRNAs
LFC

**Figure S2.**

**A)** Gene-level volcano plots of whole-genome CRISPR/Cas9 KO screens comparing wild-type (WT) to *JAK2* SK-MEL-2 cells treated with IFN-β (400 U/ml). Data represent the average of two independent screens performed on separate days. Significant hits are highlighted (*P*-adjusted <0.05, FDR <0.1).

**B)** Summary of genes conferring significant sensitivity and resistance to IFN-γ and IFN-β cytokines in SK-MEL-2 cells (*P*-adjusted value <0.05 and an average effect size of log2 fold-change > 1.5, or <-1.5 in both *JAK1* KO cell clone screens).

**C)** CRISPR/Cas9 KO screens identify genes conferring resistance or sensitivity to IFN-γ in SK-MEL-2, HT-29 and A375 cancer cell models. Scatter plots comparing log 2-fold-change values for WT control versus IFN-γ-treated conditions for each cell model. *CHD1* and *MAP3K7* are highlighted in red. Color indicates the *P* adjusted value for HT-29 or SK-MEL-2 screens, and point size indicates FDR for HT-29 or SK-MEL-2 screens. Data represent the average of two independent screens performed on separate days.

**D)** Shared and private modulators of resistance and sensitivity to IFN-γ across cancer cell lines. WT CRISPR/Cas9 KO screens in the presence or absence of IFN-γ in HT-29, SK-MEL-2 and A375. Genes with an effect size of log2 fold-change >1 or < -1 and *P* adjusted of < 0.05 are shown for two or more cell models. Total number of genes in each category is denoted by *n*. Data represent the average of two independent screens performed on separate days and are representative of two independent screens.

**E)** Gene ontology and pathway analysis of shared sensitizing and resistance hits to IFN-γ reveal cell line autonomous responses. Analysis was conducted using g:Profiler to conduct functional profiling (g:GOSt) of shared sensitizing and resistance genes (*n* = 237) between all three cell lines. Sources include; Gene Ontology Biological Process (GO:BP), Molecular Function (GO:MF), Kyoto Encyclopedia of Genes and Genomes (KEGG), Reactome pathway database (REAC).

**F)** CRISPR KO screens determine gene essentiality in HT-29 cells in the presence and absence of IFN-γ. Data is representative of gene log2 fold-changes from two independent screens performed on separate days. Results were generated using MAGeCK.

**G)** Guide RNA (gRNA) level analysis of CRISPR screens in WT and *JAK1* KO clones. Log2 fold-change (LFC) in HT-29 of each of the two gene-targeting gRNAs are highlighted in red for the two independent screens, or black for control gRNAs. All other library gRNAs are in grey.

### Figure S3

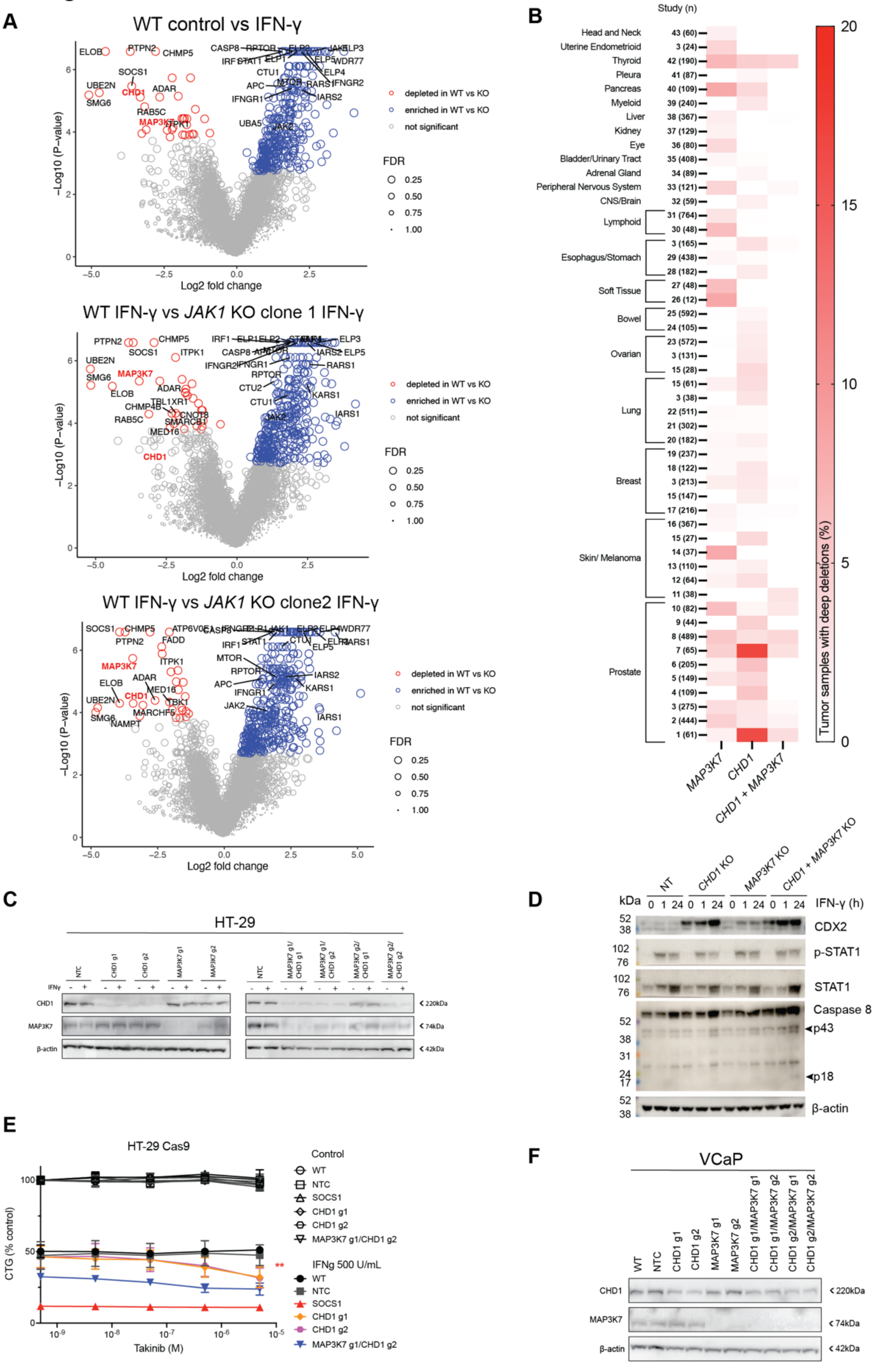
A
WT control vs IFN-γ
WT IFN-γ vs JAK1 KO clone 1 IFN-γ
WT IFN-γ vs JAK1 KO clone2 IFN-γ
−Log10 (P−value)
Log2 fold change
depleted in WT vs KO
enriched in WT vs KO
not significant
FDR
0.25
0.50
0.75
1.00
B
Study (n)
Head and Neck
Uterine Endometrioid
Thyroid
Pleura
Pancreas
Myeloid
Liver
Kidney
Eye
Bladder/Urinary Tract
Adrenal Gland
Peripheral Nervous System
CNS/Brain
Lymphoid
Esophagus/Stomach
Soft Tissue
Bowel
Ovarian
Lung
Breast
Skin/ Melanoma
Prostate
MAP3K7
CHD1
CHD1 + MAP3K7
Tumor samples with deep deletions (%)
C
HT-29
D
kDa
NT
CHD1 KO
MAP3K7 KO
CHD1 + MAP3K7 KO
IFN-γ (h)
CDX2
p-STAT1
STAT1
Caspase 8
p43
p18
β-actin
E
HT-29 Cas9
Control
WT
NTC
SOCS1
CHD1 g1
CHD1 g2
MAP3K7 g1/CHD1 g2
IFNg 500 U/mL
CTG (% control)
Takinib (M)
F
VCaP
CHD1
MAP3K7
β-actin

**Figure S3.**

**A)** Gene-level volcano plots of whole-genome CRISPR/Cas9 KO screens comparing WT cells to *JAK1* KO clones treated with IFN-γ. Data represent the average of two independent screens performed on separate days. Results were generated using MAGeCK. Significant hits are highlighted (P-adjusted <0.05, FDR <0.1).

**B)** Heatmap displaying the frequency of patient tumor samples harboring deep deletions of *CHD1* alone, *MAP3K7* alone, or both genes grouped by tumor type and study. The percentage of tumor samples with deep deletions per study from The Cancer Genome Atlas (TCGA) and other published sources are shown and listed in Supplementary Table 4. *n* denotes the total number of samples per study.

**C)** Confirmation of *CHD1* and *MAP3K7* KO in single KO and dKO cells. Western blotting of HT-29 cells in the presence or absence of IFN-γ (500 U/mL) for 1 h before analysis.

**D)** Confirmation of functional JAK-STAT signaling and increased CDX2 expression and caspase 8 cleavage in single KO and dKO in HT-29 cells. Western blotting of HT-29 cells in the presence or absence of IFN-γ (500 U/mL) for 1 h or 24 h before analysis.

**E)** IFN-γ sensitivity of *CHD1* KO and *CHD1* and *MAP3K7* dKO HT-29 cells under pharmacological inhibition of *MAP3K7* gene product TAK1. Cells were untreated or treated with IFN-γ (500 U/mL) and Takinib (5 nM - 50 µM) for six days, and cell viability was measured using Cell Titer-Glo (CTG). Data was normalized to untreated samples and represent the mean ± S.D. of three biological replicates from two independent experiments.

**F)** Confirmation of reduced *CHD1* and *MAP3K7* expression in KO and dKO cells. Western blotting analysis of VCaP cells representative of two independent experiments.

### Figure S4

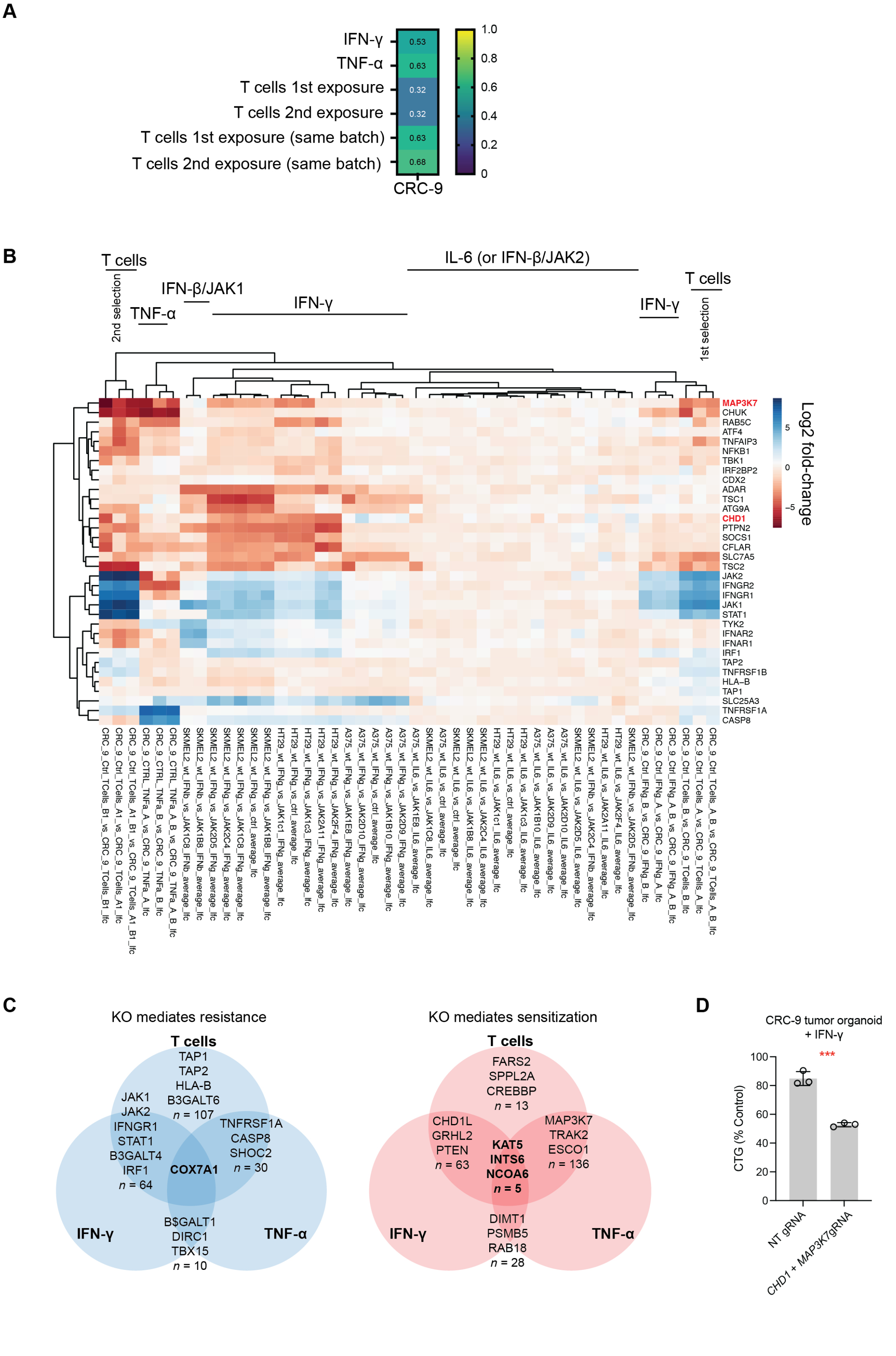
A
IFN-γ
TNF-α
T cells 1st exposure
T cells 2nd exposure
T cells 1st exposure (same batch)
T cells 2nd exposure (same batch)
0.53
0.63
0.32
0.32
0.63
0.68
CRC-9
1.0
0.8
0.6
0.4
0.2
0
B
T cells
2nd selection
TNF-α
IFN-β/JAK1
IFN-γ
IL-6 (or IFN-β/JAK2)
IFN-γ
T cells
1st selection
MAP3K7
CHUK
RAB5C
ATF4
TNFAIP3
NFKB1
TBK1
IRF2BP2
CDX2
ADAR
TSC1
ATG9A
CHD1
PTPN2
SOCS1
CFLAR
SLC7A5
TSC2
JAK2
IFNGR2
IFNGR1
JAK1
STAT1
TYK2
IFNAR2
IFNAR1
IRF1
TAP2
TNFRSF1B
HLA–B
TAP1
SLC25A3
TNFRSF1A
CASP8
Log2 fold-change
5
0
−5
C
KO mediates resistance
T cells
TAP1
TAP2
HLA-B
B3GALT6
n = 107
JAK1
JAK2
IFNGR1
STAT1
B3GALT4
IRF1
n = 64
TNFRSF1A
CASP8
SHOC2
n = 30
COX7A1
B$GALT1
DIRC1
TBX15
n = 10
IFN-γ
TNF-α
KO mediates sensitization
T cells
FARS2
SPPL2A
CREBBP
n = 13
CHD1L
GRHL2
PTEN
n = 63
KAT5
INTS6
NCOA6
n = 5
MAP3K7
TRAK2
ESCO1
n = 136
DIMT1
PSMB5
RAB18
n = 28
IFN-γ
TNF-α
D
CRC-9 tumor organoid
+ IFN-γ
***
CTG (% Control)
100
80
60
40
20
0
NT gRNA
CHD1 + MAP3K7gRNA

**Figure S4.**

**A)** Heatmap representation of replicate correlation from genome-wide tumoroid CRISPR/Cas9 KO screens in CRC-9 in the presence of different cytokines or autologous human T cells. Pearson correlation coefficient (*r*) of the log2 fold-change against the control arm is shown and results were generated using MAGeCK.

**B)** Heatmap and clustering of samples based on CRISPR/Cas9 KO screen log2 fold-changes across cell models and immunological selection pressures. Highlighted genes include *CHD1*, *MAP3K7* and other hits discussed in the main text. Each column represents a different CRISPR screen against the control sample (e.g. control versus IFN-γ, or WT versus *JAK1* KO in the presence of IFN-γ).

**C)** Venn diagram of the overlap between cytokine and T cell hits in CRC-9 tumoroid CRISPR/Cas9 KO screens. Significant gene hits are shown (*P*-adjusted < 0.05 and FDR < 0.2) with an effect size of log2 fold-change >0.5 or < -0.5 in at least two cytokine screens (IFN-γ and TNF-$\alpha$), T cells alone, or T cells and at least one cytokine screen.

**D)** Sensitivity of *CHD1* and *MAP3K7* dKO CRC-9 tumor organoids to IFN-γ. Organoids were grown in the presence or absence of IFN-γ (4000 U/mL) for six days and cell viability was measured using Cell Titer-Glo (CTG). Data was normalized to untreated samples and represent the mean ± S.D. of three biological replicates from three independent experiments.

**Figure S5**

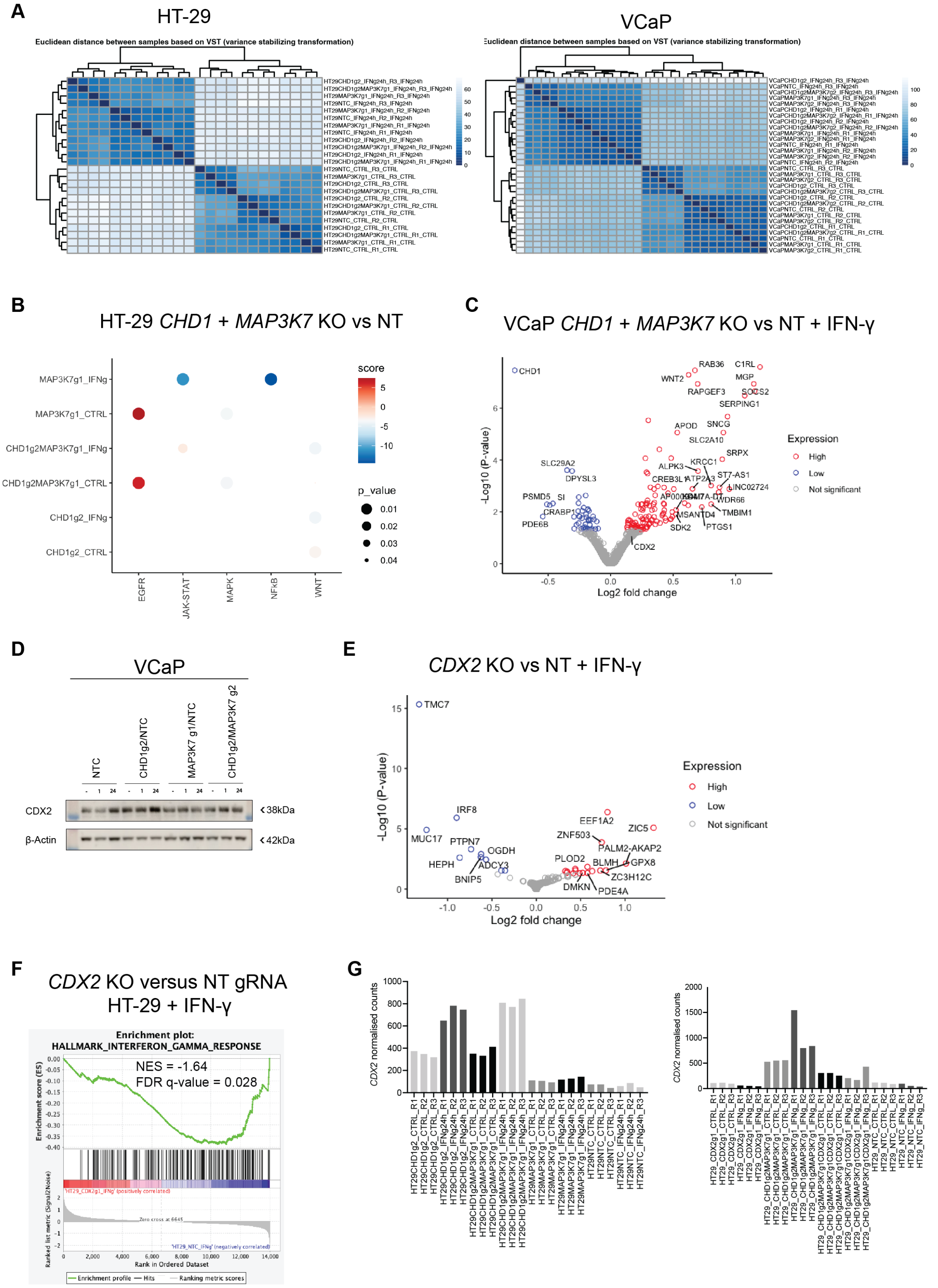
A
HT-29
Euclidean distance between samples based on VST (variance stabilizing transformation)
VCaP
B
HT-29 CHD1 + MAP3K7 KO vs NT
score
p_value
MAP3K7g1_IFNg
MAP3K7g1_CTRL
CHD1g2MAP3K7g1_IFNg
CHD1g2MAP3K7g1_CTRL
CHD1g2_IFNg
CHD1g2_CTRL
EGFR
JAK-STAT
MAPK
NFkB
WNT
C
VCaP CHD1 + MAP3K7 KO vs NT + IFN-γ
-Log10 (P-value)
Log2 fold change
Expression
High
Low
Not significant
CHD1
CDX2
D
VCaP
NTC
CHD1g2/NTC
MAP3K7 g1/NTC
CHD1g2/MAP3K7 g2
CDX2
β-Actin
38kDa
42kDa
E
CDX2 KO vs NT + IFN-γ
-Log10 (P-value)
Log2 fold change
Expression
High
Low
Not significant
TMC7
IRF8
MUC17
EEF1A2
ZIC5
F
CDX2 KO versus NT gRNA
HT-29 + IFN-γ
Enrichment plot:
HALLMARK_INTERFERON_GAMMA_RESPONSE
NES = -1.64
FDR q-value = 0.028
Enrichment score (ES)
Rank in Ordered Dataset
Enrichment profile
Hits
Ranking metric scores
G
CDX2 normalised counts

**Figure S5.**

**A)** RNA-seq sample heatmap demonstrating clustering from HT-29 or VCaP cells based on hclust Euclidean distance of normalized RNA counts from DESeq2. All independent replicates were performed on separate days and had a Pearson correlation coefficient >0.9, except for VCaP CHD1g2 IFNg24h replicate 3 (R3), which clustered separately and was excluded from downstream analysis (Methods).

**B)** Pathway analysis reveals NF-$\kappa$B and JAK-STAT signaling in HT-29 cells. Pathway activity for each single and dKO cell line in the presence or absence of IFN-γ (500 U/mL) for 24 h. P values are derived from decoupleR PROGENy scores. PROGENy pathways without significant changes in any cell line are not plotted.

**C)** Differential gene expression comparing dKO and NT gRNA harboring VCaP cells cultured in IFN-γ (500 U/mL) for 24 h. Data represents the average of three independent biological replicates, highlighting differentially expressed transcripts with a P-adjusted value of <0.05.

**D)** Confirmation of *CHD1* and *MAP3K7* KO and upregulation of *CDX2* in dKO VCaP cells. Western blotting of VCaP cells in the presence or absence of IFN-γ (500 U/mL) for 1 or 24 h before analysis.

**E)** *CDX2* KO causes downregulation of *IRF8* in HT-29 cells. Differential gene expression comparing *CDX2* KO and NT gRNA harboring HT-29 cells cultured in IFN-γ (500 U/mL) for 72 h. Data represents the average of three independent biological replicates, highlighting differentially expressed transcripts with a P-adjusted value of <0.05.

**F)** The IFN-γ pathway is downregulated in *CDX2* KO cells. Gene-set enrichment analysis for the Hallmark IFN-γ signature between *CDX2* KO and NT gRNA harboring HT-29 cells cultured in IFN-γ (500 U/mL) for 72 h. Normalized enrichment score (NES) and false discovery rate (FDR) q-value of correlation are shown.

**G)** *CDX2* normalized counts across replicates from RNA-seq data in HT-29 cells. Normalized RNA counts were determined using DESeq2.

Figure S6

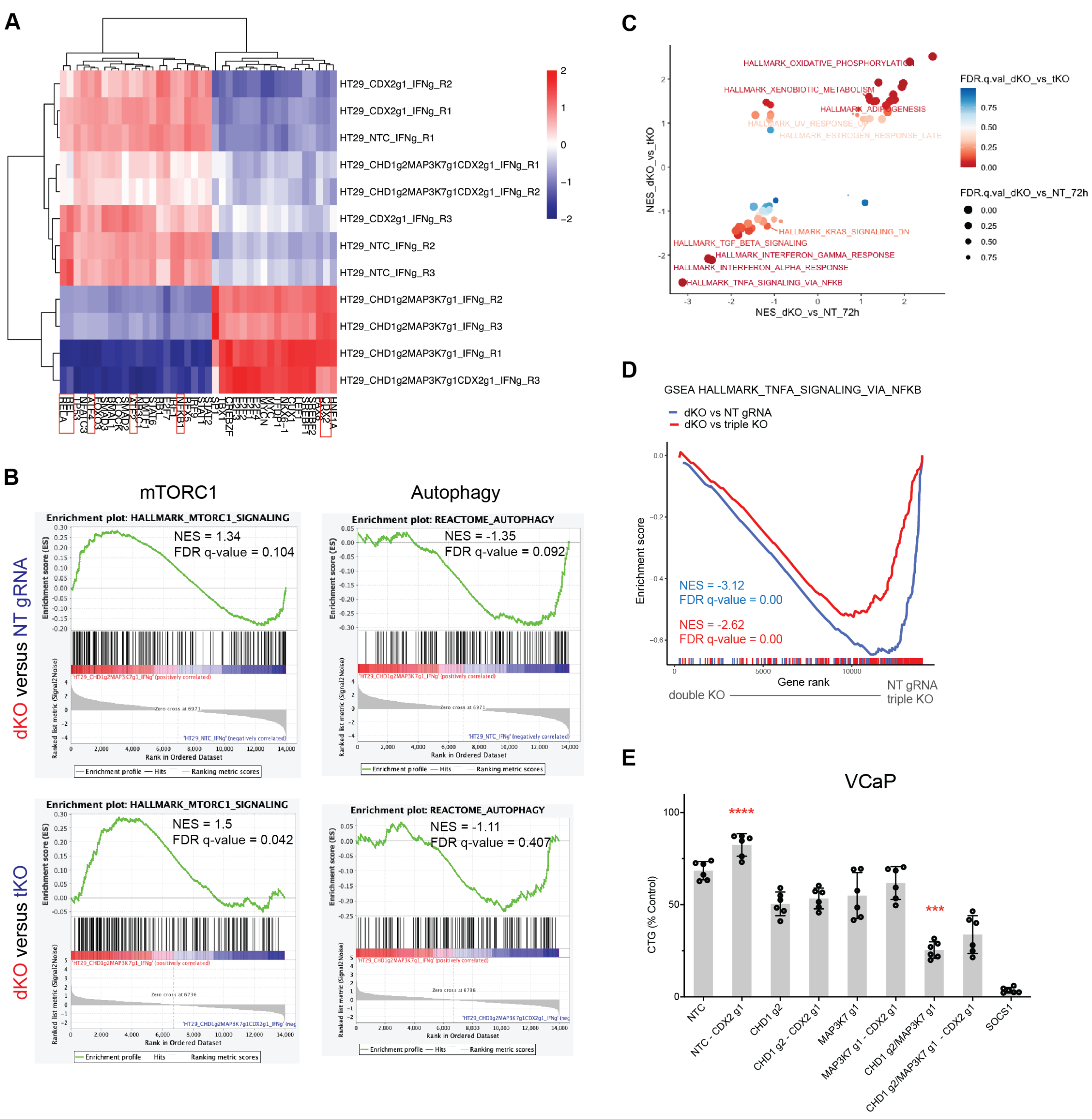


**Figure S6.**

**A)** RNA sequencing analysis reveals differential NF-κB, autophagy and IFN-γ linked transcription factor activity in *CDX2, CHD1* and *MAP3K7* KO cells in response to IFN-γ. Heatmap and hierarchical clustering of transcription factor activity in *CDX2* KO, dKO and tKO HT-29 cells in the presence of IFN-γ (500 U/mL) for 72 h. Data was analyzed with decoupleR using DESeq2 log-transformed counts. Transcription factors of interest highlighted in red boxes.

**B)** *CDX2* KO reverses the enhanced mTORC and autophagy expression signatures of dKO cells in response to IFN-γ. Gene-set enrichment analysis of hallmark mTORC1 and reactome autophagy signature between dKO, tKO and NT gRNA harboring HT-29 cells cultured in IFN-γ (500 U/mL) for 72 h. Normalized enrichment score (NES) and false discovery rate (FDR) q-value of correlation are shown.

**C)** *CDX2* KO reverses dKO induced gene programs. Gene-set enrichment analysis signatures of gene sets in HT-29 cells. Comparisons are between dKO vs NT and dKO vs tKO in the presence of IFN-γ (500 U/mL) for 72 h. Normalized enrichment score (NES) and false discovery rate (FDR) q-value of correlation are shown.

**D)** *CDX2* KO rescues the reduced TNF-$\alpha$ signaling via NF-$\kappa$B in dKO cells. Gene-set enrichment analysis of hallmark TNF-$\alpha$ via NF-$\kappa$B signature in dKO, tKO, and NT gRNA harboring HT-29 cells cultured in IFN-γ (500 U/mL) for 72 h. Normalized enrichment score (NES) and false discovery rate (FDR) q-value of correlations are shown.

**E)** Sensitivity of KO, dKO, tKO, or *SOCS1* KO VCaP cells to IFN-γ. Cells were untreated or treated with IFN-γ (500 U/mL) and cell viability was measured using Cell Titer-Glo (CTG). Data was normalized to untreated samples and represent the mean ± S.D of two independent experiments, each performed in technical triplicate. Two-tailed, unpaired Student's t-test; *** = P <0.001, **** = P <0.0001, not significant = n.s.

### Figure S7

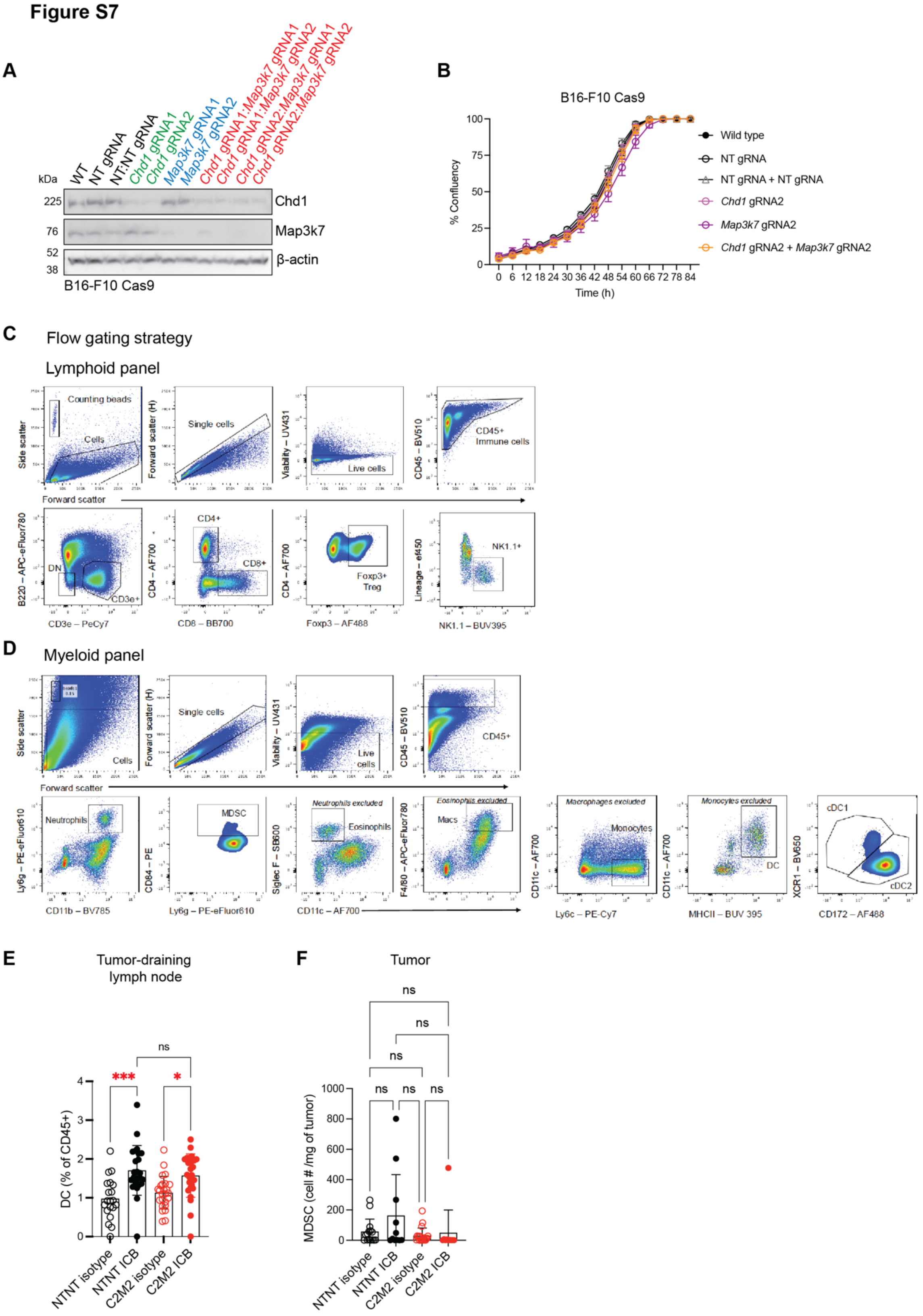
A
WT
NT gRNA
NT:NT gRNA
Chd1 gRNA1
Chd1 gRNA2
Map3k7 gRNA1
Map3k7 gRNA2
Chd1 gRNA1:Map3k7 gRNA1
Chd1 gRNA1:Map3k7 gRNA2
Chd1 gRNA2:Map3k7 gRNA1
Chd1 gRNA2:Map3k7 gRNA2
kDa
225
76
52
38
Chd1
Map3k7
β-actin
B16-F10 Cas9
B
B16-F10 Cas9
% Confluency
Time (h)
Wild type
NT gRNA
NT gRNA + NT gRNA
Chd1 gRNA2
Map3k7 gRNA2
Chd1 gRNA2 + Map3k7 gRNA2
C
Flow gating strategy
Lymphoid panel
Counting beads
Cells
Side scatter
Forward scatter
Single cells
Forward scatter (H)
Viability – UV431
Live cells
CD45 – BV510
CD45+ Immune cells
B220 – APC-eFluor780
DN
CD3e+
CD3e – PeCy7
CD4+
CD8+
CD4 – AF700
CD8 – BB700
Foxp3+ Treg
Foxp3 – AF488
Lineage – ef450
NK1.1+
NK1.1 – BUV395
D
Myeloid panel
CD45+
Neutrophils
Ly6g – PE-eFluor610
CD11b – BV785
MDSC
CD84 – PE
Neutrophils excluded
Eosinophils
Siglec F – SB600
CD11c – AF700
Eosinophils excluded
Macs
F4/80 – APC-eFluor780
Macrophages excluded
Monocytes
Ly6c – PE-Cy7
Monocytes excluded
DC
MHCII – BUV 395
cDC1
cDC2
XCR1 – BV650
CD172 – AF488
E
Tumor-draining lymph node
DC (% of CD45+)
***
*
ns
NTNT isotype
NTNT ICB
C2M2 isotype
C2M2 ICB
F
Tumor
MDSC (cell # /mg of tumor)
ns
NTNT isotype
NTNT ICB
C2M2 isotype
C2M2 ICB

**Figure S7.**

**A)** Confirmation of *Chd1* and *Map3k7* KO in single KO and *Chd1/Map3k7* in dKO B16-F10 cells. Western blotting of B16-F10 Cas9 cells.

**B)** KO of *Chd1*, *Map3k7*, or both (dKO), does not affect cell growth in B16-F10 cells. Cell proliferation was monitored using an Incucyte. Data represent the mean ± S.D. of three technical replicates and are representative of two independent experiments performed on separate days.

**C)** Progressive flow cytometry gating strategy for immunophenotyping of lymphoid cells in mouse syngeneic, subcutaneous B16-F10 tumors, lymph nodes and spleen. For lineage gating we used CD172α, Siglec-F, XCR1, CD64, CD11b, I-AE/I-E, CD11c, F4/80, Ly6G, Ly-6C markers. Counting beads were used to measure absolute cell counts.

**D)** Progressive flow cytometry gating strategy for immunophenotyping of myeloid cells in mouse syngeneic, subcutaneous B16-F10 tumors, lymph nodes and spleen. For lineage gating we used CD3, NK1.1, CD5, CD19, B220 markers. Counting beads were used to measure absolute cell counts.

**E)** Confirmation of dendritic cell (DC) influx into inguinal tumor draining lymph nodes (tdLNs) following systemic immune checkpoint blockade (ICB). Flow cytometry analysis of DCs in tdLNs in the presence of anti-PD-1 and anti-CTLA-4 (ICB) or isotype (Iso.) control. Data represents the mean ± S.D of three independent experiments. One-way ANOVA, * $P = 0.0259$; ***$P = 0.0002$. NT NT Iso. $n$=20. NT NT ICB $n$=24. dKO Iso. $n$=25. dKO ICB $n$=25.

**F)** There is no difference in Myeloid derived suppressor cells (MDSC) within NT control and *Chd1* and *Map3k7* dKO tumors. Flow cytometry analysis of tumor MDSCs in the presence of either anti-PD-1 and anti-CTLA-4 immune checkpoint blockade (ICB) or isotype (Iso) control. Data represents the mean ± S.D of three independent experiments. N.s; not significant. One-way ANOVA. NT NT Iso. $n$=15. NT NT ICB $n$=11. dKO Iso. $n$=19. dKO ICB $n$=10.

**Figure S8**

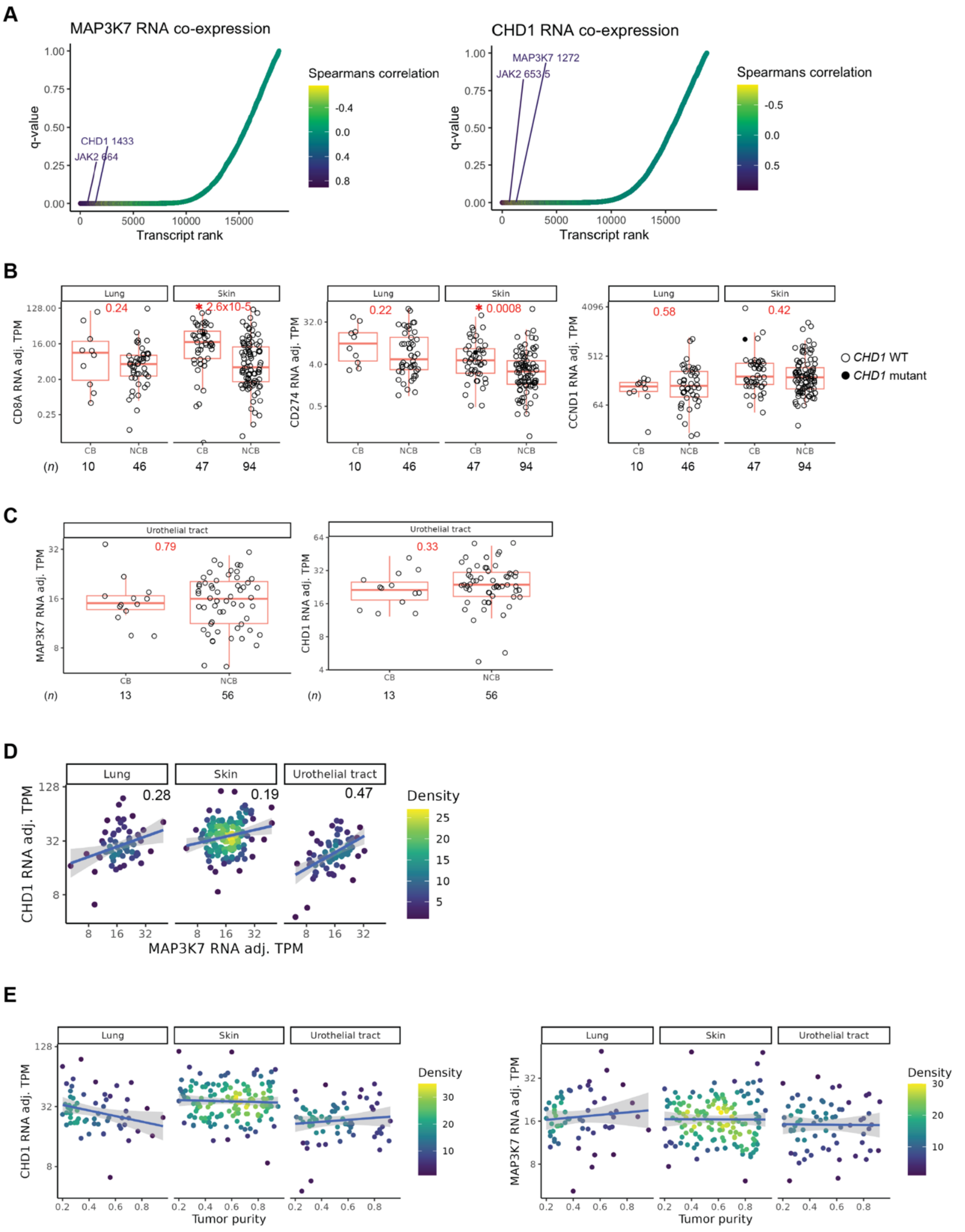


**Figure S8.**

**A)** *CHD1* and *JAK2* RNA expression are co-correlated with the expression of *MAP3K7*. Co-expression of genes with *MAP3K7* in prostate cancer patients. *CHD1*, *P* = 7.19e-22 and *MAP3K7*, *P* = 2.47e-20.

**B)** Box-plot displaying mRNA expression of tumor *CD8A*, *CD274* (PD-L1) and *CCND1* expression (adjusted transcripts per million; adj. TPM) and clinical responses to ICB in lung cancers and melanoma from the Hartwig Medical Foundation[90]. Clinical benefit, CB; no clinical benefit, NCB. Significance was assessed using the Wilcoxon signed-rank test and *n* denotes the number of patients. Boxplots represent the median, interquartile range (IQR) and whiskers are the lowest and highest values within 1.5 × IQR.

**C)** Box-plot displaying mRNA expression of tumor *CHD1* and *MAP3K7* expression (adjusted transcripts per million; adj. TPM) and clinical responses to ICB in urothelial carcinomas from the Hartwig Medical Foundation[90]. Clinical benefit, CB; no clinical benefit, NCB. Significance was assessed using the Wilcoxon signed-rank test and *n* denotes the number of patients. Boxplots represent the median, interquartile range (IQR) and whiskers are the lowest and highest values within 1.5 × IQR.

**D)** *CHD1* and *MAP3K7* RNA expression is correlated in ICB treated tumor sample data. RNA expression of *CHD1* and *MAP3K7* in ICB treated lung, skin and urothelial tract tumor patients from the HMF. RNA expression is measured as transcripts per million (TPM). R is a measure of Pearson's correlation. The shaded region represents the 95 % confidence interval.

**E)** *CHD1* and *MAP3K7* expression are not correlated with tumor sample purity. RNA expression and tumor purity estimates of *CHD1* and *MAP3K7* in ICB-treated lung, skin and urothelial tract tumors from the HMF. RNA expression is measured as transcripts per million (TPM). The shaded region represents the 95 % confidence interval.
