## Supplemental Notes for "CRISPR screens in the context of immune selection identify *CHD1* and *MAP3K7* as mediators of cancer immunotherapy resistance"

**Supplemental Note 1**

Independent tumoroid screens with different batches of T cell cultures resulted in lower levels of screen replicate correlation (*r* = 0.31-0.32) than cytokine selection screens (*r* = 0.63-0.53), likely due to different levels of tumor cell killing between T cell batches (Fig. S4C-D). However, two rounds of tumoroid selection with the same T cell preparations generated more consistent data (*r* = 0.68-0.63; Fig. S4E), with increasing effect-sizes from the first to the second round of cytotoxic selection with T cells (Fig. S4B).

**Supplemental Note 2**

Transcriptional signatures of increased androgen receptor signaling following loss of either or both genes (Fig. 3B) could explain why they are most frequently mutated in prostate cancer and associated with resistance to anti-androgen treatment^1^. Recent studies further support the unanticipated link between cancer cell JAK-STAT and androgen receptor signaling^2,3^.

**Supplemental Note 3**

Despite previous reports suggesting *Chd1* loss in tumors can affect myeloid-derived suppressor cell (MDSC) recruitment^4^, we did not observe a change in MDSC frequencies (Fig. S8F), possibly due to the low overall abundance of MDSCs in these B16-F10 tumors.

**Supplemental Note 4**

We identify a conserved role for amino acid sensing and mTOR signaling in mediating cancer cell sensitivity to cytokines. mTOR inhibits autophagy, providing a functional link to a known immune evasion pathway^5^ involved in the lysosomal destruction of pro-death complexes formed in response to cytotoxic cytokines^6^. These results imply that rapamycin could exert part of its immunosuppressive effect by acting directly on tissues, rather than the immune cell compartment alone^7^.

**Supplemental Note 5**

In lung cancers, *MAP3K7* expression was more predictive of ICB response than *CD8A* or *CD274* (PD-L1) (Fig. S8B) and a known mediator of ICB resistance, *CCND1*^8^, however this trend was not significant in urothelial tumors (Fig. S8C). *CHD1* and *MAP3K7* expression were modestly correlated in tumor samples from patients treated with ICB (*r* = 0.47-0.19; Fig. S8D), but expression was not correlated with tumor purity, implying this association is independent of tumor immune cell content (Fig. S8E). *CDX2* tumor expression was not significantly correlated with ICB outcome (p >0.14, Wilcoxon signed-rank test).
